# The ETS Transcription Factor ERF controls the exit from the naïve pluripotent state

**DOI:** 10.1101/2021.02.01.429223

**Authors:** M. Vega-Sendino, T. Olbrich, D. Tillo, A. D. Tran, C. N. Domingo, M. Franco, P. C. FitzGerald, M. J. Kruhlak, S. Ruiz

**Author notes:** Corresponding author, @sruizmacias.

## Abstract

The naïve epiblast undergoes a transition to a pluripotent primed state during embryo implantation. Despite the relevance of the FGF pathway during this period, little is known about the downstream effectors regulating this signaling. Here, we examined the molecular mechanisms coordinating the naïve to primed transition by using inducible ESC to genetically eliminate all RAS proteins. We show that differentiated RAS^KO^ ESC remain trapped in an intermediate state of pluripotency with naïve-associated features. Elimination of the transcription factor ERF overcomes the developmental blockage of RAS-deficient cells by naïve enhancer decommissioning. Mechanistically, ERF regulates NANOG expression and ensures naïve pluripotency by strengthening naïve transcription factor binding at ESC enhancers. Moreover, ERF negatively regulates the expression of the *de novo* methyltransferase DNMT3B, which participates in the extinction of the naïve transcriptional program. Collectively, we demonstrated an essential role for ERF controlling the exit from naïve pluripotency during the progression to primed pluripotency.

**Teaser:** ERF is the MAPK-dependent switch controlling the transition between naïve and primed pluripotency during embryonic development.

## INTRODUCTION

Embryonic cells residing within the inner cell mass (ICM) are pluripotent as they have the potential to generate all cell lineages of the organism. The state of pluripotency found in the preimplantation epiblast, prior to any lineage specification, is usually defined as naïve (*1*). However, as the embryo expands and develops after implantation, epiblast cells become individually fated, although still retain pluripotent features. This state of pluripotency associated to the post-implantation epiblast is defined as primed (*2, 3*). More than distinct pluripotent states, naïve and primed can be viewed as different phases of a coordinated developmental progression where naïve unbiased cells differentiate responding to inductive cues to initiate a multi-lineage decision commitment at gastrulation (*2, 3*).

Naïve and primed pluripotent states can be captured *in vitro* by defined culture conditions. Mouse embryonic stem cells (ESC) can be propagated in naïve conditions by using a combination of a MEK inhibitor (PD0325091) plus a glycogen synthase kinase-3 inhibitor (CHIR99021; hereafter referred as the 2i condition) (*4*). ESC grown under these conditions resemble embryonic cells residing in the pre-implantation embryonic (E) post-fertilization day E3.5-E4.25 embryo and have been extensively used to study the ground state of pluripotency. In addition, cultures of embryonic cells that retain primed pluripotency can also be established from post-implantation embryos (*5*). These epiblast stem cells (EpiSC) resemble transcriptionally the late epiblast layer of the post-implantation embryo at E6.0-E6.5. Both types of cells, naïve and primed, share the expression of core pluripotent transcription factors (TF), such as OCT4 and SOX2. However, 2i-ESC express specific naïve TF (REX1, NANOG or KLF4), which are absent in EpiSC, whereas EpiSC express epiblast specific genes (OCT6 or OTX2), absent in ESC. Importantly, these pluripotent cell types are interconvertible by modifying the culture conditions or expressing specific TF (*3*). Recently, a distinct intermediate state of pluripotency has been identified, the rosette pluripotent state (*6*). Rosette embryonic stem cells (RSC) co-express naïve TF and the primed marker OTX2. Inhibition of WNT signaling drives the pluripotent transition from naïve to the rosette state while further activation of the RAS/MAPK signaling promotes the progression to primed pluripotency (*6*). Furthermore, additional states of intermediate pluripotency characterized by germ cell specification capacity have also been described, formative stem cells (FSC) and chimera pluripotent stem cells (XPSC) (*7, 8*). These intermediates rely on exogenous (XPSC) or autocrine (FSC) FGF signaling for their self-renewal in contrast to the requirements for RSC. In summary, pluripotency can be considered as a dynamic property associated to different stem cell states supported by defined pluripotent transcriptional networks.

Exit from naïve pluripotency during implantation is essential to fate epiblast cells residing in the ICM with inductive signals prior to gastrulation. However, the molecular mechanisms coordinating the naïve to primed transition are not fully understood. Moreover, despite the relevance of the RAS pathway during this critical period, little is known about the downstream effectors regulating the MAPK transcriptional program. We recently identified the transcriptional repressor ERF, a member of the E26 transformation specific (ETS) family, as an important regulator downstream of the RAS pathway in ESC (*9*). ERF is a TF that shuffles between the nuclei and cytoplasm in ESC in a phosphorylation-dependent manner (*10*). In the absence of RAS/MAPK signaling, ERF remains unphosphorylated and chromatin-bound in the nuclei while growth factor stimulation keeps ERF inactive in the cytoplasm by ERK-dependent phosphorylation (*10–12*). Importantly, the precise role of ERF during early embryonic development and pluripotent transitions is unclear. Here, we show that ERF plays a dual role in the transition from naïve to primed pluripotency. First, ERF binds to ESC super-enhancers to ensure optimal expression of naïve TF in the absence of FGF signaling. Second, activation of MAPK signals induces the release of chromatin-bound ERF, an event that is necessary and sufficient to trigger full commitment to primed pluripotency. In addition, we also show that ERF controls the expression of the *de novo* methyltransferase DNMT3B through LIN28 regulation, leading to the inactivation of the naïve transcriptional program. Our data demonstrate that ERF is the MAPK-dependent switch that controls the progression to primed pluripotency.

## RESULTS

### ERF is expressed in the naïve pluripotent epiblast

We first sought to determine the precise expression timing for ERF during embryonic development. For this, we collected mouse embryos at different developmental stages and performed immunofluorescence analyses. We determined that ERF expression peaks around E3.5-E4.0 and is coincidental with the naïve pluripotent epiblast (Fig. 1A-C). ERF is expressed in the naïve inner cell mass (ICM) and co-expressed with the naïve pluripotent markers NANOG and KLF4 (Fig. 1B, C). In addition, we also detected ERF in the trophectoderm (TE), consistent with its reported requirement for chorionic trophoblast differentiation (Fig. 1B, C) (*13*). Exit from naïve pluripotency *in vivo* and downregulation of naïve associated markers strongly correlated with decreased expression of ERF (Fig. 1B, C). We confirmed this expression pattern by analyzing single-cell RNAseq datasets from embryos at different stages (fig. S1A) (*14*). These results showed that ERF is upregulated during ICM formation and it is quickly downregulated before implantation, suggesting a role for ERF in the naïve epiblast.

**Fig. 1.**
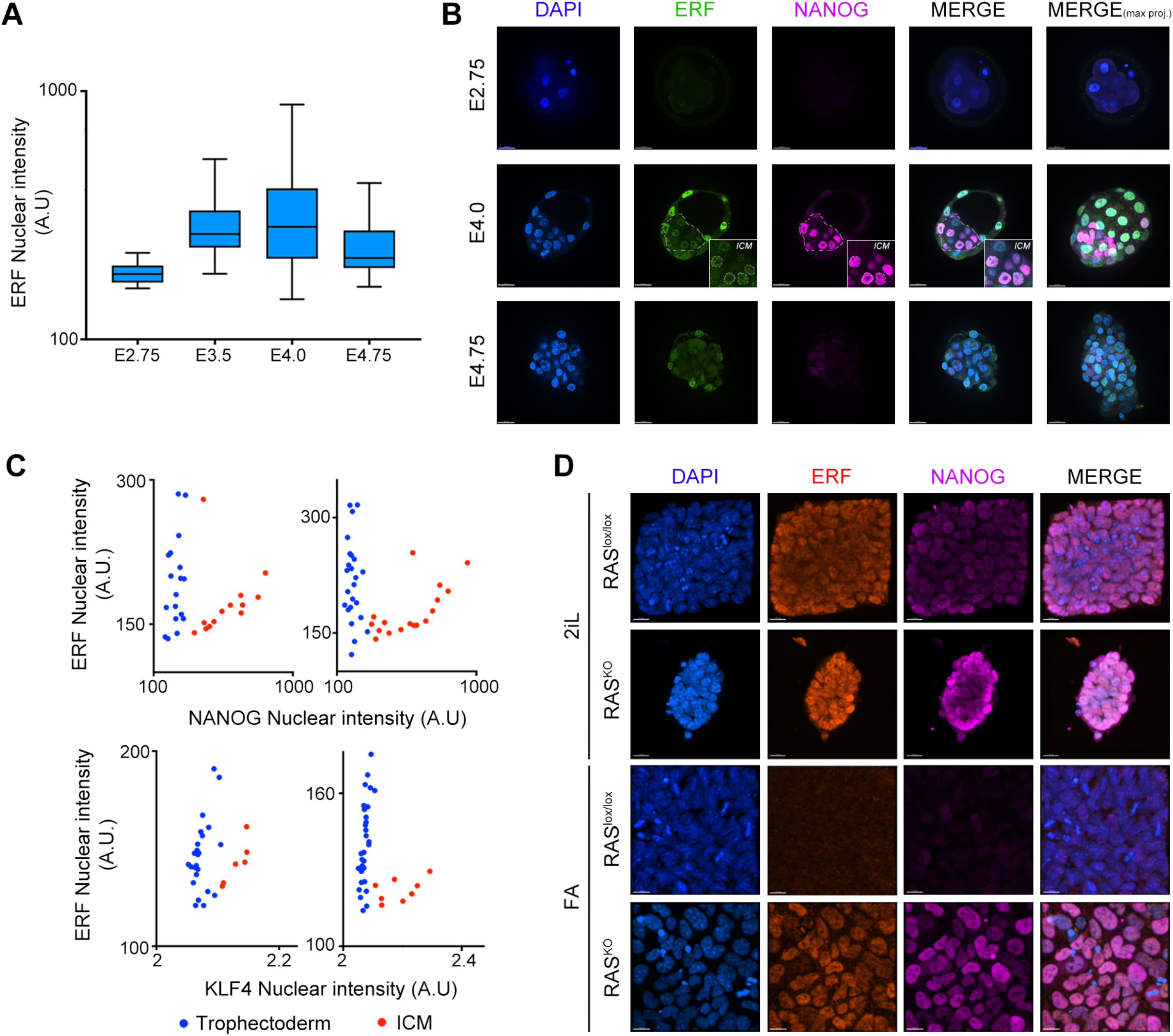
ERF expression correlates with naïve pluripotency markers. (**A**) Graph showing mean nuclear fluorescent intensity for ERF in mouse embryos at different days of embryonic development (E). (**B**) Immunofluorescence analysis of NANOG and ERF in mouse embryos at E2.75, E4.0 and E4.75. Note that ERF is expressed in both ICM and TE. However, at E4.75 ERF and NANOG are mostly downregulated in epiblast cells. Dashed line highlights the ICM. DAPI was used to visualize nuclei. Scale bars, 20μm. (**C**) Graphs showing relative nuclear fluorescence intensity of ERF and NANOG (upper plots) and ERF and KLF4 (lower plots). Every dot represents one single nucleus, and each plot corresponds to an individual E3.5 embryo. Two representative examples are shown but at least 10 embryos were analyzed. (**D**) Immunofluorescence analysis of 2iL and FA treated RAS^lox/lox^ and RAS^KO^ ESC and stained for ERF (red) and NANOG (purple). DAPI was used to visualize nuclei. Scale bars, 15μm.

To validate our observations and assess the role of the MAPK pathway in ERF levels, we used an ESC model deficient in H-RAS and N-RAS with a conditional knockout K-RAS triggered by the addition of 4-hydroxytamoxifen (OHT) (RAS^lox/lox^ hereafter) (*9, 15*). These ESC allowed us to generate cells devoid of all RAS proteins. We next applied a well-established *in vitro* protocol using FGF2 and Activin-A (FA, hereafter) to differentiate naïve ESC, growing in 2i+LIF (2iL) conditions, to post-implantation epiblast like stem cells (EpiLSC) and examined ERF levels (*16, 17*). While RAS^lox/lox^ 2iL-ESC showed high levels of ERF, these cells exhibited negligible levels when differentiated into EpiLSC (Fig. 1D). ERF is quickly phosphorylated upon differentiation and the decrease in protein levels is due to transcriptional repression (fig. S1B) (*18*). We next examined whether this downregulation depended on FGF/MAPK activation. Indeed, RAS-deficient cells (RAS^KO^) retained elevated levels of ERF after differentiation (Fig. 1D). Together, these results pointed to a role for ERF in the exit from naïve pluripotency in a MAPK-dependent manner.

### Downregulation of ERF is necessary for the successful exit from naïve pluripotency

To evaluate the implication of ERF in the exit from naïve pluripotency, we examined its expression along with markers of naïve pluripotency during the formation of embryonic rosettes. In this assay, single cell suspensions of ESC embedded in Matrigel exit from naïve pluripotency and develop into polarized rosettes that undergo lumenogenesis, mimicking the morphogenic events of the epiblast (*19*). This *in vitro* system recapitulates the development of the ICM during per-iimplantation. Importantly, ESC rosettes generated under 2iL conditions lack lumen, continue to express naïve markers, and eventually become disorganized over time (*20*). This showed that exit from naïve pluripotency is necessary for successful polarization and lumenogenesis. Upon 2iL removal and resuspension in Matrigel, RAS^lox/lox^ rosettes showed polarization and lumen, expression of the sialomucin protein podocalyxin (PDX) and downregulation of the naïve marker NANOG (Fig. 2A-C and fig. S2A and B). However, RAS^KO^ rosettes developed into a disorganized group of cells, which lacked expression of PDX and retained naïve pluripotency markers (Fig. 2A-C and fig. S2A and B). To test whether ERF regulates the exit from naïve pluripotency, we generated ERF-knockouts (ERF^KO^) and evaluated rosette formation in all different genotypes (RAS^lox/lox^, ERF^KO^, RAS^KO^, and RAS^KO^; ERF^KO^). Elimination of ERF is sufficient to induce exit from naïve pluripotency and to rescue the failed morphogenic events (polarization and lumenogenesis) observed in RAS^KO^ rosettes (Fig. 2A-C and fig. S2A and B). To further support these observations, we generated reporter cell lines in our ESC by replacing the endogenous coding sequence of the gene REX1 (also known as ZFP42) with a short half-life form of eGFP (REX1-deGFP). Similar REX1-deGFP reporter ESC have been widely used as a faithful system to monitor exit from naïve pluripotency (*21*). Induction of EpiLSC by FA in RAS^lox/lox^ and ERF^KO^ ESC demonstrated efficient downregulation of the reporter while RAS^KO^ ESC showed no signs of downregulation (Fig. 2D). However, exit from naïve pluripotency was efficiently achieved in RAS^KO^; ERF^KO^ ESCs (Fig. 2D). Finally, we also examined the clonogenicity potential of RAS^KO^; ERF^KO^ ESC primed to exit naïve pluripotency. For this, cultures of ESC (RAS^lox/lox^, ERF^KO^, RAS^KO^, and RAS^KO^; ERF^KO^) were withdrawn of 2iL for 2 days and plated back in 2iL conditions for a colony forming assay. Cells that exit naïve pluripotency are irreversibly committed and have lost the ability to generate colonies in 2iL conditions. Indeed, while RAS^KO^ ESC were able to generate alkaline phosphatase positive colonies and remained trapped in a naïve pluripotent state, RAS^KO^; ERF^KO^ ESCs lost this ability (Fig. 2E). Our results demonstrate a predominant role for ERF controlling the exit from naïve pluripotency.

**Fig. 2.**
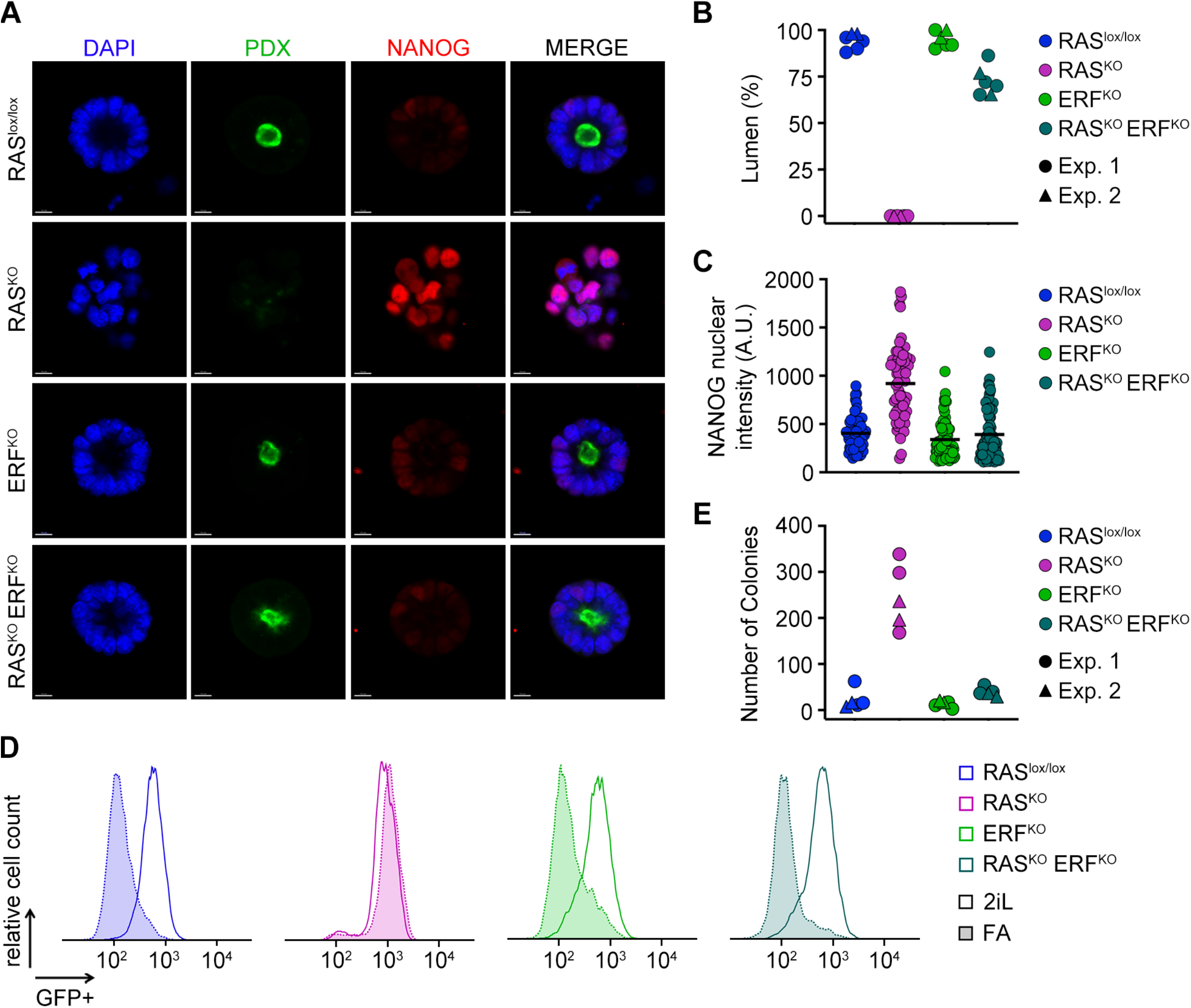
Successful exit from naïve pluripotency requires downregulation of ERF. (**A**) Central confocal optical sections of RAS^lox/lox^, ERF^KO^, RAS^KO^, and RAS^KO^; ERF^KO^ embryonic cell rosettes embedded in matrigel 48 hours after seeding and stained for NANOG (red) and podocalyxin (PDX; green). DAPI was used to visualize nuclei. Scale bars, 10μm. (**B**) Graph showing the percentage of embryonic rosettes generating a lumen (PDX+) in all genotypes. Two independent experiments are shown and at least, a total of 50 rosettes were counted per sample. (**C**) Graph showing mean NANOG nuclear fluorescence intensity per nuclei in embryonic rosettes 48 hours after seeding. One representative experiment is shown and a total of 70 nuclei from different rosettes were counted per sample. (**D**) Flow cytometry analysis of REX1-deGFP reporter ESC from all genotypes in 2iL-ESC and EpiLSC after 48 hours of induction with FA. Three independent experiments were performed but one representative experiment is shown. (**E**) Graph showing the number of alkaline phosphatase positive colonies in a colony forming assay using ESC from all genotypes. Two independent experiments are shown with at least two technical replicates.

### ERF controls the progression to primed pluripotency

To obtain mechanistic insights on the rescue mediated by the loss of ERF during the exit from naïve pluripotency, we performed RNA sequencing (RNAseq) analysis of ESC cultured under naïve conditions (2iL) or differentiated into EpiLSC in FA. Principal component plot (PCA) analysis segregated the samples based on their differentiation status alongside PC1, and MAPK activity alongside PC2 (Fig. 3A). Interestingly, FA-RAS^KO^ ESC are localized at an intermediate state between naïve and primed pluripotency (Fig. 3A). FA-RAS^KO^ ESC are characterized by the intermediate to high expression of naïve pluripotent markers as well as of primed associated genes including OTX2 (Fig. 3B, C). Indeed, while naïve pluripotent ESC are characterized by a NANOG^+^/OTX2^-^/OCT6^-^ state, FA-RAS^KO^ ESC showed a NANOG^+^/OTX2^+^/OCT6^-^ state (Fig. 3B, C). Furthermore, although FA-RAS^lox/lox^ ESC have undetectable levels of ERF upon differentiation, FA-RAS^KO^ ESC retain high ERF expression (Fig. 3C). Using available datasets that evaluate temporal transcriptional dynamics during the transition from naïve to primed pluripotency, we determined that FA-RAS^KO^ ESC transcriptionally resembled to ESC primed for 12-24 hours (Fig. 3D and fig. S3A) (*18*). Interestingly, FA-RAS^KO^ ESC are reminiscent of the recently described intermediate pluripotent states, rosette and formative pluripotent states (RSC, FSC and XPSC) (*6–8*). PCA analyses showed that PC1 (differentiation status) placed all intermediate pluripotent states in a similar transcriptional space but PC2 segregated FA-RAS^KO^ ESC from rosette or formative pluripotent states (Fig. 3E). Hierarchical clustering analysis based on the expression of naïve and primed markers revealed that RSC and FA-RAS^KO^ ESC are transcriptionally more comparable, likely to the defective MAPK signaling in both intermediate states (Fig. 3F). Finally, we observed that prolonged culture of FA-RAS^KO^ ESC in FGF2/Activin-A/XAV939 (FAX) conditions demonstrated their trapping in this intermediate state of pluripotency (fig. S3B). Our results showed that loss of ERF is necessary and sufficient to overcome the developmental blockage of FA-RAS^KO^ ESC in its intermediate pluripotent state. Indeed, RNAseq data revealed that loss of ERF in RAS^KO^; ERF^KO^ ESC restored the overall gene expression profile to be indistinguishable from FA-RAS^lox/lox^. In summary, our data point to ERF as the MAPK-dependent switch that triggers full commitment to primed pluripotency.

**Fig. 3.**
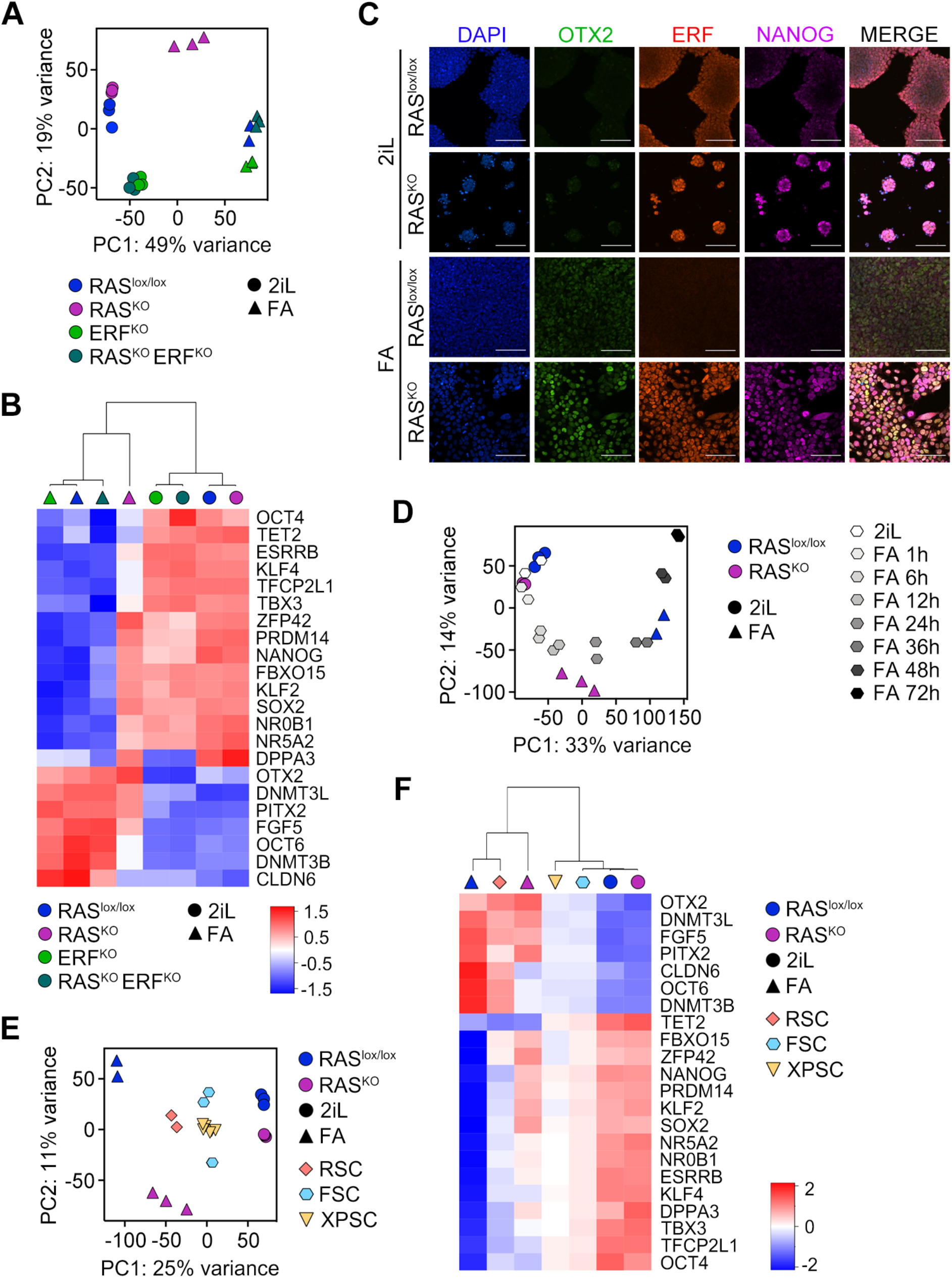
ERF controls the transition to primed pluripotency. (**A**) PCA plot of RNAseq data of RAS^lox/lox^, ERF^KO^, RAS^KO^, and RAS^KO^; ERF^KO^ ESC cultured in naïve conditions (2iL) or induced to differentiate to EpiLSC (FA) for 48 hours. Three replicates per condition are shown. (**B**) Heatmap generated from RNAseq data from samples described in A) showing the average from three replicates. (**C**) Immunofluorescence analysis of 2iL and FA treated RAS^lox/lox^ and RAS^KO^ ESC stained for OTX2 (green), ERF (red) and NANOG (purple). DAPI was used to visualize nuclei. Scale bars, 100μm. (**D**) PCA plot of RNAseq datasets (three replicates) showing 2iL and FA treated RAS^lox/lox^ and RAS^KO^ ESC along with RNAseq datasets (two replicates) from a time course experiment during EpiLSC induction (0, 1, 6, 12, 24, 36, 48 and 72 hours) (*18*). (**E**) PCA plot of RNAseq datasets showing 2iL and FA treated RAS^lox/lox^ and RAS^KO^ ESC along with RNAseq datasets from RSC (ESC lines grown in LIM (LIF, IWP2 (WNT inhibitor) and MEKi), FSC and XPSC (*6–8*). At least two replicates are shown. (**F**) Heatmap generated from RNAseq data from samples described in (E) showing the average from two or three replicates as applicable to each sample.

### Chromatin-bound ERF ensures an optimal naïve pluripotency state

The expression of ERF correlates with high expression levels of naïve markers *in vivo* and *in vitro* and is quickly downregulated upon induction to primed pluripotency (Fig. 1). Moreover, ERF is enriched in a total of 2074 bona-fide ESC enhancers identified in RAS^KO^ ESC (Supplemental Data S1 and 2) (*9*). Among these, ERF binds to most of the super-enhancers identified in ESC (198/231) (Supplemental Data S3) (*9, 22, 23*). ERF-bound super-enhancers are associated to highly transcribed naïve genes (KLF4, ESRRB, PRDM14, NANOG, TBX3 or ZFP42) (some examples in fig. S4A) as well as to general pluripotent genes (OCT4 or SOX2). Although ERF is considered to be a transcriptional repressor, we hypothesized that ERF might play a different role at ESC enhancers. To explore the relevance of ERF at these enhancers, we first examined the level of occupancy of the pluripotent transcription factors OCT4, SOX2 and NANOG (O, S and N, respectively) in the subset of ERF-bound 2074 ESC enhancers. We observed increased occupancy of bound OSN at these compared to a non-ERF bound randomized set of different 2074 enhancers (Fig. 4A). In addition, ERF-bound ESC enhancers are also characterized by higher H3K27Ac levels, increased p300 binding and chromatin accessibility detected by ATACseq (fig. S4B). These results suggested that ERF could be regulating the activity of these enhancers, and thus, the expression of their associated genes. Therefore, we examined the expression levels of essential naïve pluripotent genes associated to ERF-bound super-enhancers in RAS^lox/lox^ and ERF^KO^ ESC grown in 2iL conditions. Interestingly, NANOG, PRDM14 and ZFP42 showed decreased expression in 2iL-ERF^KO^ ESC, while TBX3, KLF4 and ESRRB do not (Fig. 4B). It has been shown that NANOG promotes chromatin accessibility and binding of pluripotent factors to enhancers (*24*). Thus, we hypothesized that reduced levels of NANOG could affect the expression of additional naïve-associated genes. Indeed, ERF^KO^ cells showed an overall decreased expression of the naïve pluripotent transcriptional network (Fig. 4C). Consistent with these results, unidimensional PCA analysis (PC1) segregating samples based on their differentiation status showed that 2iL-ERF^KO^ ESC are biased toward differentiation (Fig. 4D). These combined results revealed that ESC growing under naïve conditions required ERF binding at enhancers to maintain an optimal naïve pluripotent state. To confirm this observation, we employed native Cut&Run sequencing to examine SOX2 and NANOG occupancy in RAS^lox/lox^ and ERF^KO^ ESC grown in 2iL conditions. As predicted, both pluripotent transcription factors showed an overall decreased enrichment in ESC enhancers in naïve ERF^KO^ ESC (Fig. 4E, F and fig. S4C). These data support our findings and revealed an unexpected unique role for ERF in promoting naïve pluripotency.

**Fig. 4.**
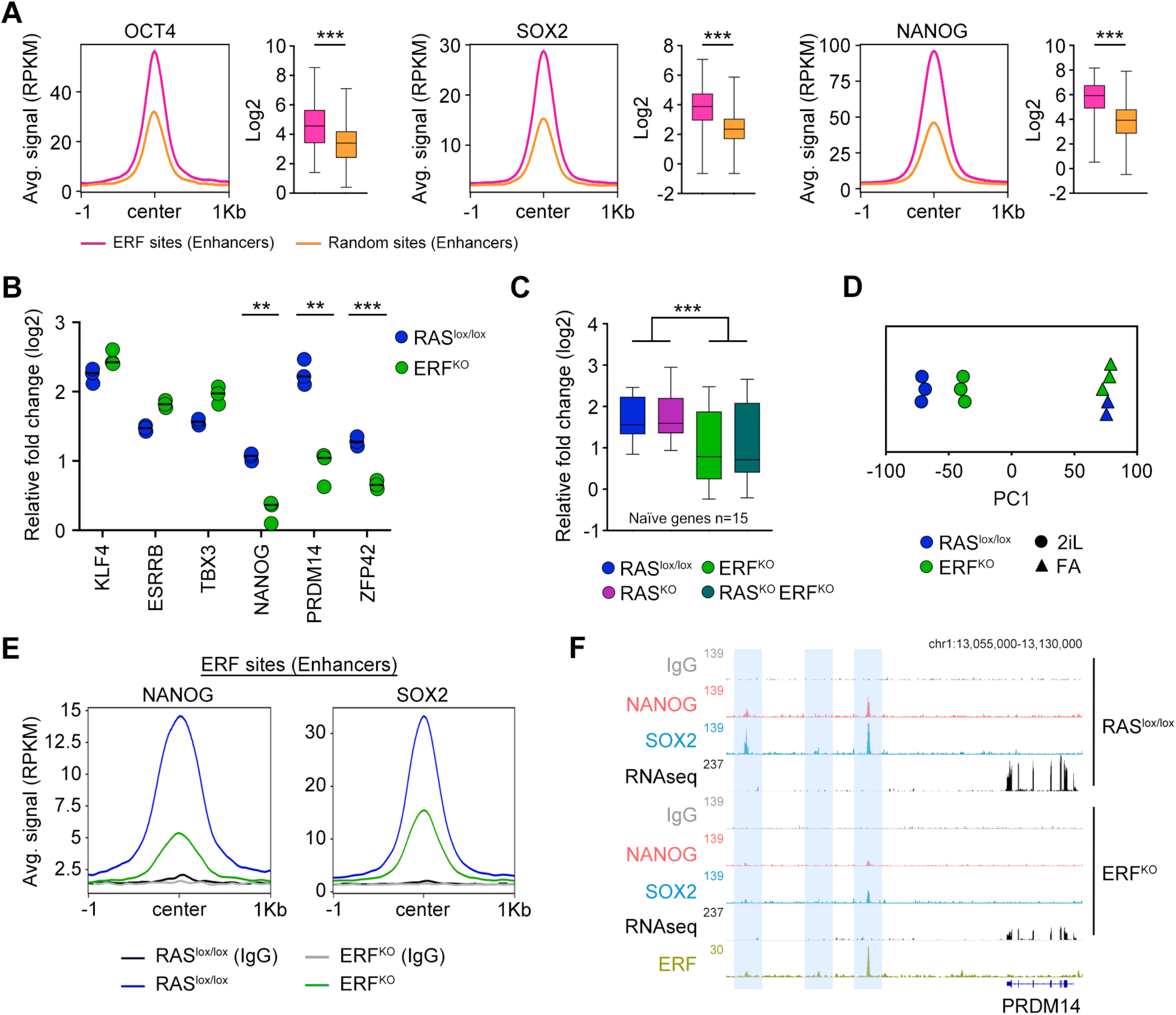
ERF ensures optimal naïve pluripotent transcription factor expression in ESC. (**A**) ChIPseq read density plot (RPKM) showing OCT4, SOX2 and NANOG occupancy at 2074 ERF-binding sites at enhancers (pink) or 2074 randomly selected non-ERF bound enhancers (orange). Graphs show quantifications of the TF enrichment in each set of sites. *** = p<0.001, T-student. Data was obtained from (*43*). (**B**) Graph showing relative fold change (log2) expression of the indicated genes in RAS^lox/lox^ and ERF^KO^ ESC grown in 2iL. Genes showing at least a 50% reduction are highlighted. For each gene, data was normalized to the average across all samples. *** = p<0.001; ** = p<0.01. T-student. Data is shown as triplicates. (**C**) Graph showing the relative fold change (log2) expression of a subset of 15 naïve associated genes (*Klf4, Esrrb, Tfcp2l1, Tbx3, Klf2, Elf3, Klf8, Nanog, Prdm14, Tcl1, Zfp42, Nrob1, Prmd16, Dazl and Crxos*) in the different genotypes in 2iL conditions. For each gene, data was normalized to the average across all samples. *** = p<0.001. T-student. (**D**) Unidimensional PCA plot of RNAseq data of RAS^lox/lox^ and ERF^KO^ ESC cultured in naïve conditions (2iL) or induced to differentiate (FA) during 48 hours to EpiLSC. Three replicates per condition are shown. PC1 separates samples by differentiation status. (**E**) Cut&Run read density plot (RPKM) showing NANOG and SOX2 occupancy in the set of 2074 ERF-binding sites at enhancers in RAS^lox/lox^ (blue) and ERF^KO^ (green) ESC cultured in 2iL. Corresponding inputs (IgG) are also shown as reference control. (**F**) Genome browser tracks showing NANOG and SOX2 occupancy and RNAseq RPKM read count at the PRDM14 gene in the indicated genotypes. ERF binding profile in RAS^KO^ ESC is also shown. Inputs (IgG) are also shown as a reference control. ERF binding sites are highlighted.

### The naïve enhancer landscape is active in FA-RAS^KO^ ESC

Inactivation of the naïve transcriptional network is necessary for the successful progression to primed pluripotency. This is associated with decommission of naïve enhancers by downregulation of naïve transcription factors including ESRRB, KLF4 or NANOG and a widespread OCT4 genomic relocation (*25*). Indeed, OCT4 shifts from enhancers associated with key players in naïve pluripotency and engages in new enhancer elements at genes implicated in post-implantation development. Using datasets for OCT4 occupancy in ESC and EpiLSC and based on the relative occupancy between both cellular states, we defined top OCT4 sites preferentially enriched in ESC (4759 sites, OCT4^ESC^), EpiLSC (2921 sites, OCT4^EpiLSC^) and commonly shared between the two (9144 sites, OCT4^Common^) (fig. S5A and Supplemental Data S4-6). Genes associated with OCT4^ESC^ sites are mostly downregulated during the transition to primed pluripotency whereas genes associated to OCT4^EpiLSC^ sites are upregulated (*25*). Out of all OCT4 bound sites, a total of 12.52% (596/4759 ERF/OCT4^ESC^ sites), 13.62% (1246/9144 ERF/OCT4^Common^ sites) and 0.15% (44/2921 ERF/OCT4^EpiLSC^ sites) were co-occupied by ERF in naïve ESC (Fig. 5A). This suggested that ERF does not play a specific role in genes associated with EpiLSC-specific sequences but instead in regulating OCT4^ESC^ and OCT4^Common^ sites. We first focused on OCT4^ESC^ sites as we observed that FA-RAS^KO^ ESC remain trapped in an intermediate state between naïve and primed pluripotency and showed expression of naïve pluripotent markers (Fig. 3A-C). Differentially expressed genes associated with ERF/OCT4^ESC^ sites were mostly downregulated during FA differentiation in RAS^lox/lox^ ESC (Fig. 5B). While many of these genes did not change their expression level in FA-RAS^KO^ ESC, those that changed showed a lower differential expression extent (Fig. 5B). Enhancer decommission of naïve enhancers by OCT4 relocation is followed by changes in enhancer chromatin patterns including decreased levels of H3K27ac (*24*). Thus, we examined whether the naïve enhancer landscape is still fully active in FA-RAS^KO^ ESC by performing native Cut&Run sequencing to evaluate the levels of H3K27ac as a marker for active enhancers and NANOG occupancy in ESC and EpiLSC from all genotypes. Interestingly, 2iL-RAS^KO^ ESC showed increased acetylation and NANOG occupancy at ERF/OCT4^ESC^ sites (Fig. 5C, D), a phenotype that is also observed in all OCT4^ESC^ sites (fig. S5B). This increase in H3K27ac levels does not result in major overall transcriptional changes and the expression level of naïve associated genes is similar between RAS^lox/lox^ and RAS^KO^ ESC (Fig. 4C). This observation supports the idea that levels of H3K27ac does not necessarily determine enhancer activity but rather discriminates between active or poised enhancers (*26*). Exit from naïve pluripotency correlated with an overall decrease in H3K27ac levels, negligible chromatin-bound NANOG and reduced expression of naive-associated genes in RAS^lox/lox^ ESC (Fig. 5C, D and fig. S5B). However, FA-RAS^KO^ ESC showed elevated gene expression, NANOG occupancy and H3K27ac levels at ERF/OCT4^ESC^ sites to, in some cases, a comparable level as detected in naïve RAS^lox/lox^ ESC (Fig. 5C, D). Together, these data showed that naïve-specific associated OCT4 binding sites remain fully active in the FA-RAS^KO^ ESC intermediate state.

**Fig. 5:**
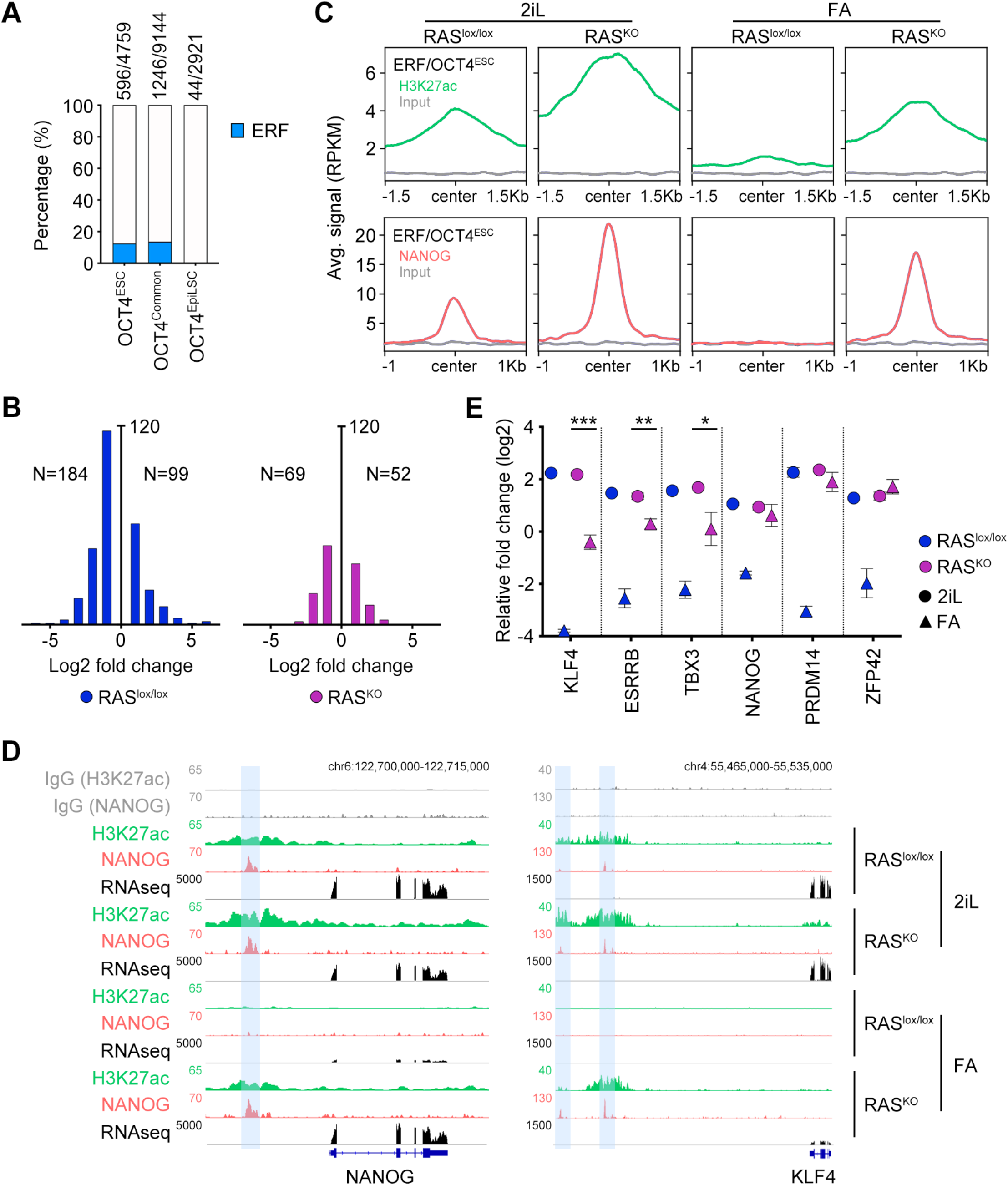
The naïve pluripotent transcription factor network is active in FA-RAS^KO^ ESC. (**A**) Plot showing the percentage of OCT4 binding sites co-occupied by ERF in ESC, EpiLSC and common between ESC and EpiLSC. (**B**) Histogram plots showing fold expression changes (log2) for genes associated to ERF/OCT4^ESC^ that are characterized by differential expression between ESC and EpiLSC in RAS^lox/lox^ and RAS^KO^ cells. N indicates the total number of genes that are up or downregulated. Genes were associated at every binding site by proximity using GREAT (ref PMID: 20436461). (**C**) Cut&Run read density plot (RPKM) showing H3K27ac (plots above, green) and NANOG (plots below, red) occupancy in ERF/OCT4^ESC^ sites in RAS^lox/lox^ and RAS^KO^ cultured in 2iL or differentiated to EpiLSC (FA). Corresponding inputs (IgG) are also shown as reference control. (**D**) Genome browser tracks showing H3K27ac deposition, NANOG occupancy and RNAseq RPKM read count at the KLF4 and NANOG genes in the indicated genotypes. Inputs (IgG) are also shown as a reference control. (**E**) Graph showing the relative fold change (log2) expression of the indicated ERF-bound super-enhancer associated genes in RAS^lox/lox^ and RAS^KO^ cultured in 2iL or differentiated to EpiLSC (FA). Genes showing at least a 50% reduction are highlighted. For each gene, data was normalized to the average across all samples. *** = p<0.001; ** = p<0.01; *=<0.05 T-student. Data shown are averages from triplicates.

We next focused specifically on the regulatory sequences and expression level of naïve transcription factors associated with ERF-bound super-enhancers. Among these, the expression level of ERF-dependent genes (NANOG, PRDM14 and ZFP42) is unaffected in FA-RAS^KO^ ESC and is comparable to that of RAS^lox/lox^ ESCs as ERF is still expressed (Fig. 5E). However, ERF-independent genes (ESRRB, TBX3 and KLF4) showed a significant decrease in expression, suggesting that additional mechanisms ensure optimal expression of these genes in 2iL-ESC (Fig. 5E). Collectively, these results demonstrated that FA-RAS^KO^ ESC retain an active naïve transcriptional network. Although some naïve markers showed a marked decreased expression at this stage, elevated levels of NANOG, PRDM14 or ZFP42 likely sustain the naïve like-state in FA-RAS^KO^ ESC mediated by ERF-dependent mechanisms.

### OTX2 co-occupy binding sites with NANOG in FA-RAS^KO^ ESC

Global reorganization of OCT4 genomic binding is driven by increased expression of OTX2 during naïve to primed transition (*25*). In fact, ectopic OTX2 expression in 2iL-ESC shows that it behaves as a pioneering factor engaging in previously inaccessible enhancer sites, relocates OCT4 to these sites, and induces the expression of primed-associated genes (*25*). OTX2 expression is independent of MEK signals but is efficiently repressed by the WNT pathway, thus explaining the elevated levels of OTX2 in rosette-like ESC (*6*). High levels of OTX2 have also been found to be associated to formative pluripotency (*7, 8*). In addition, we also detected similar high OTX2 levels in FA-RAS^KO^ ESC compared to FA-RAS^lox/lox^ (Fig. 6A). We have shown that ERF binding sustains optimal naïve transcription factor expression (Fig. 4) and thus, we asked whether ERF is involved in OTX2 regulation. Indeed, ERF binds to the super-enhancer region associated with OTX2 in RAS^KO^ ESC. Furthermore, the low levels of OTX2 expression observed in 2iL-RAS^lox/lox^ ESC are further decreased in 2iL-ERF^KO^ ESC (Fig. 6A, B). This decrease correlates with lower NANOG and SOX2 binding and suggests that ERF binding might prevent further OTX2 repression in the absence of FGF signaling (Fig. 6B). In agreement, FA-RAS^KO^ ESC showed strong NANOG enrichment as well as OTX2 itself, which could potentially sustain its own expression after NANOG downregulation (fig. S6A). We next focused on the ERF/OCT4^Common^ sites as they gained OTX2 and H3K27ac enrichment in EpiLSC compared to ESC (*24*). Differentially expressed genes associated with ERF/OCT4^Common^ sites were mostly upregulated in FA-RAS^lox/lox^ (Fig. 6C). Similar to what we observed with genes associated with ERF/OCT4^ESC^ sites, many of these genes did not change their expression levels in FA-RAS^KO^ ESC and those that changed showed a lower differential expression extent (Fig. 6C). Based on the elevated levels of NANOG expression in FA-RAS^KO^ ESC, we hypothesized that ERF/OCT4^Common^ sites also retained chromatin bound NANOG. While FA-RAS^lox/lox^ showed negligible enrichment of NANOG at these sites, FA-RAS^KO^ ESC retained NANOG bound (Fig. 6D). In addition, we also observed stronger OTX2 enrichment at these sites in FA-RAS^KO^ ESC compared to FA-RAS^lox/lox^ (Fig. 6D). Interestingly, we also detected OTX2 binding in FA-RAS^KO^ ESC at the ERF/OCT4^ESC^ sites, naïve associated sequences that are decommissioned during the transition to EpiLSC (Fig. 6D). These observations were also confirmed globally in all OCT4 sites (fig. S6B). Furthermore, OTX2 as well as NANOG were also strongly enriched at the OCT4^EpiLSC^ sites in FA-RAS^KO^ ESC, suggesting that, besides OCT4, OTX2 might also relocate additional naïve pluripotent factors (if expressed) to these sites prior to full differentiation (Fig. 6D and fig. S6B). Finally, we examined whether OTX2 and NANOG enrichment correlated with increased expression of post-implantation epiblast genes associated with ERF/OCT4^Common^ sites in FA-RAS^KO^ ESC. We found genes showing a similar level of expression in FA-RAS^KO^ ESC compared to FA-RAS^lox/lox^ (Fig. 6E and fig. S6C). Our data demonstrate that FA-RAS^KO^ ESC show promiscuous OTX2 and NANOG binding at OCT4^ESC^ sites and OCT4^EpiLSC^ sites, which correlates with the expression of both naïve and primed markers in preparation for the transition towards primed pluripotency.

**Fig. 6.**
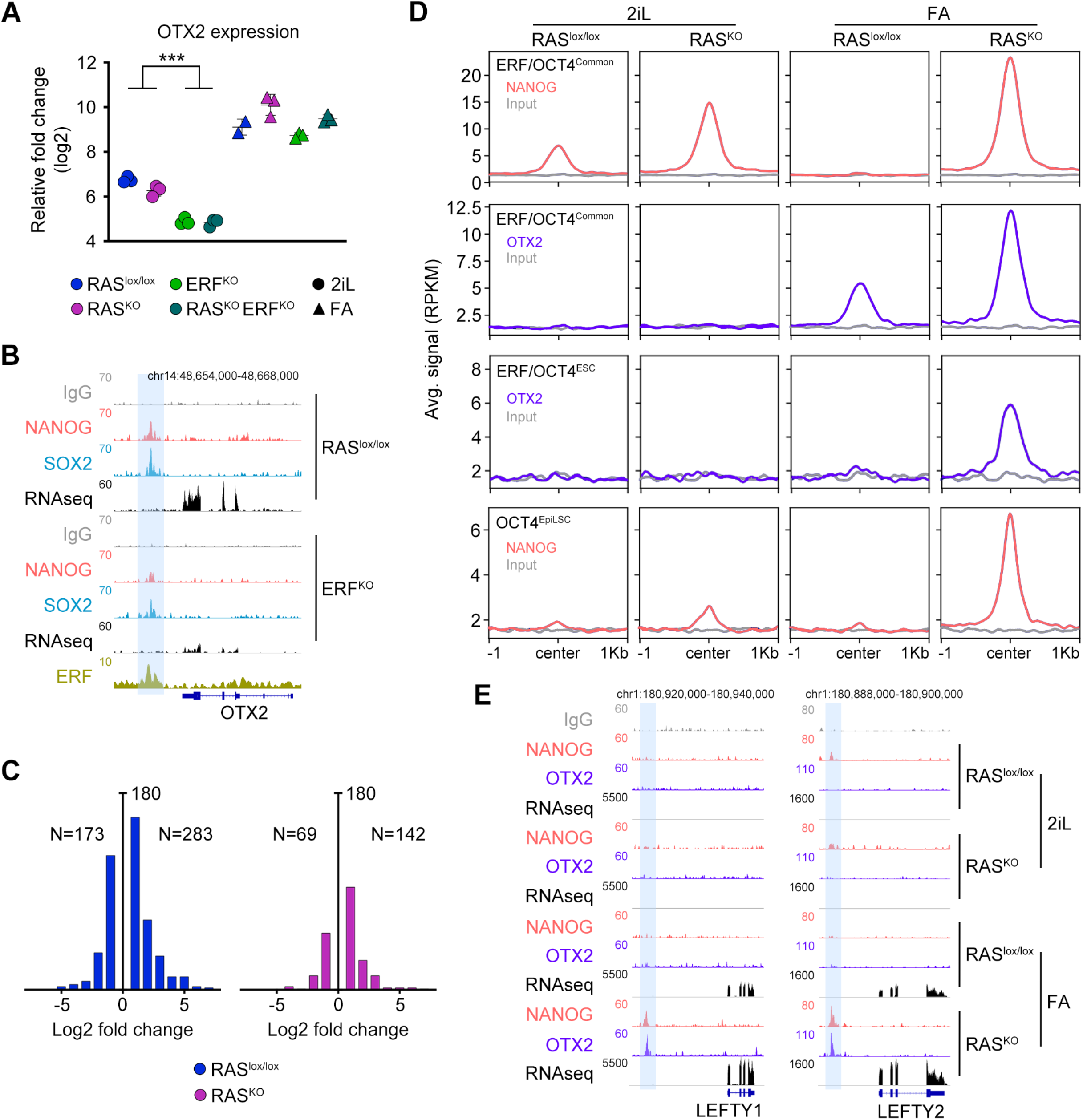
OTX2 shows promiscuous binding in naïve and primed genes in FA-RAS^KO^ ESC. (**A**) Plot showing the relative fold change (log2) expression for OTX2 in ESC from all genotypes cultured in 2iL or differentiated to EpiLSC (FA). Data was normalized to the average across all samples. ***= p<0.001; T-student. Data shown from triplicates. (**B**) Genome browser tracks showing SOX2 and NANOG occupancy and RNAseq RPKM read count at the OTX2 gene in RAS^lox/lox^ and ERF^KO^ ESC cultured in 2iL. ERF binding profile in RAS^KO^ ESC is also shown. Inputs (IgG) are also shown as a reference control. ERF binding sites are highlighted. (**C**) Histogram plots showing fold expression changes (log2) for genes associated to ERF/OCT4^Common^ that showed differential expression between ESC and EpiLSC in RAS^lox/lox^ and RAS^KO^ cells. N indicates the total number of genes that are up or downregulated. Genes were associated at every binding site by proximity using GREAT (ref PMID: 20436461). (**D**) Cut&Run read density plot (RPKM) showing NANOG (red) and OTX2 (purple) occupancy in the indicated ERF/OCT4^Common^, ERF/OCT4^ESC^ and ERF/OCT4^EpiLSC^ sites in RAS^lox/lox^ and RAS^KO^ ESC cultured in 2iL or differentiated to EpiLSC (FA). Corresponding inputs (IgG) are also shown as reference control. (**E**) Genome browser tracks showing NANOG and OTX2 occupancy and RNAseq RPKM read count at the LEFTY1 and LEFTY2 genes in RAS^lox/lox^ and RAS^KO^ ESC cultured in 2iL or differentiated to EpiLSC (FA). Inputs (IgG) are also shown as a reference control. Blue squares showed ERF binding sites.

### ERF controls the expression of LIN28 proteins

We next investigated how ERF controls the exit from the developmental blockage of RAS-deficient cells by mining our RNAseq data (Fig. 3). By combining differential gene expression between our different genetic conditions and differentiation status together with nearby ERF binding occupancy, we identified LIN28A and B as putative regulators. Expression of LIN28A/B is low in 2iL-ESC but is upregulated by FGF signaling during the transition to primed pluripotency. Of note, LIN28A and/or LIN28B-deficient ESC showed impaired conversion into EpiLSC, revealing a critical role for these proteins in regulating the exit from naïve pluripotency (*27*). Induction of EpiLSC differentiation in RAS^lox/lox^ and ERF^KO^ ESC demonstrated efficient LIN28s upregulation while FA-RAS^KO^ ESC showed low or negligible levels of LIN28A and LIN28B, respectively (Fig. 7A). ERF deficiency restored the levels of LIN28 proteins in FA-RAS^KO^; ERF^KO^ (Fig. 7A). Interestingly, 2iL-ERF^KO^ ESC showed already elevated levels of LIN28 proteins before differentiation, suggesting that ERF mediates a direct negative regulation on these genes. Indeed, we found ERF bound to an enhancer region in the second intron of LIN28 proteins in RAS^KO^ ESC (Fig. 7B and fig. S7A). Moreover, transition to EpiLSC correlated with increased levels of H3K27ac at the LIN28 enhancers, and thus, LIN28 expression (Fig. 7B).

**Fig. 7.**
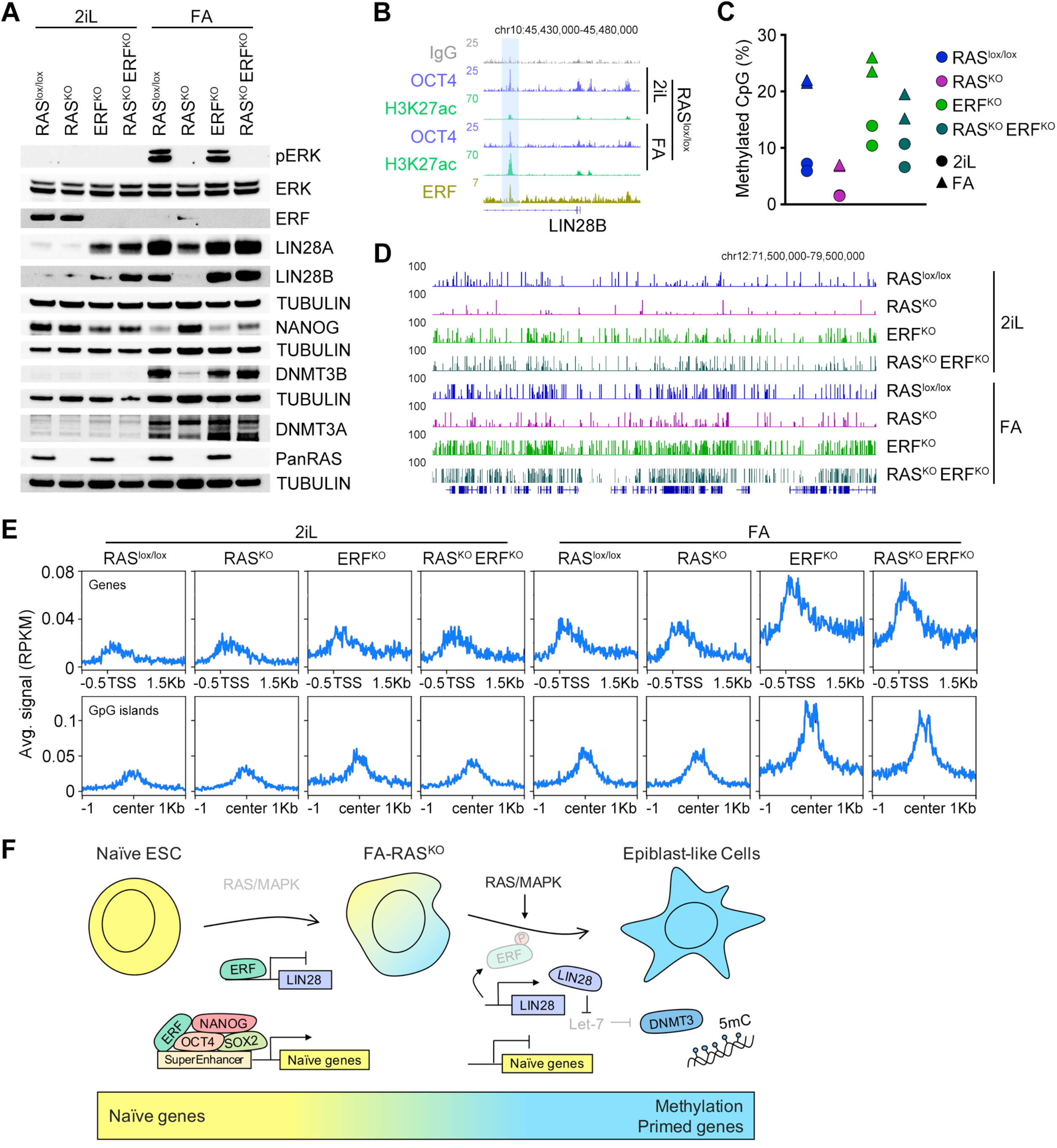
ERF controls the levels of *de novo* methylation during transition to EpiLSC. (**A**) Western blot analysis of the indicated proteins performed in ESC from all genotypes cultured in 2iL or differentiated to EpiLSC (FA). Tubulin levels for the corresponding blots are shown as a loading control. One representative experiment is shown but at least two independent clones per condition and genotype were used. (**B**) Genome browser tracks showing H3K27ac deposition and OCT4 occupancy at the LIN28B gene in RAS^lox/lox^ ESC cultured in 2iL or differentiated to EpiLSC (FA). ERF binding profile in RAS^KO^ ESC is also shown. Inputs (IgG) are also shown as a reference control. ERF binding sites are highlighted. (**C**) Graph showing the percentage of methylated CpG sites identified by RRBS in ESC from all genotypes cultured in 2iL or differentiated to EpiLSC (FA). (**D**) Genome browser tracks showing as a representative example the level of methylation at CpG sites in a region of chromosome 12 in ESC from all genotypes cultured in 2iL or differentiated to EpiLSC (FA). (**E**) %CpG methylation averaged across the TSS of all protein-coding mouse genes (upper panels) or centered at CpG islands (lower panels) in ESC from all genotypes cultured in 2iL or differentiated to EpiLSC (FA). (**F**) Schematic model showing the dual role of ERF during the naïve to primed pluripotent transition. In the absence of FGF signaling, ERF ensures an optimal level of expression for naïve transcription factors. Following ERF phosphorylation and gene silencing, ESC are able to exit and transition into EpiLSC along with global CpG methylation and silencing of the naïve transcriptional network.

LIN28 proteins are RNA binding proteins known for binding to and inactivating the let-7 microRNA family. Functionally, let-7 microRNAs target a number of mRNA transcripts for degradation including MYC, RAS, HMGA2 and the two *de novo* methyltransferases DNMT3A and DNMT3B (*28*). Indeed, naïve ESC exhibit low levels of genome wide CpG methylation, which increase during the transition to EpiLSC, correlating with the silencing of the naïve transcriptional program (*29, 30*). Accordingly, induction of RAS^lox/lox^ and ERF^KO^ ESC into EpiLSC by FA demonstrated efficient upregulation of DNMT3 proteins while FA-RAS^KO^ ESC showed low levels of DNMT3B (Fig. 7A). As expected, expression of DNMT3 proteins was efficiently rescued in FA-RAS^KO^; ERF^KO^ (Fig. 7A). We next examined the levels of 5-methylcytosine (5mC) by dot blot in all different genotypes, RAS^lox/lox^, ERF^KO^, RAS^KO^, and RAS^KO^; ERF^KO^, grown under naïve or primed conditions (fig. S7B). The low levels of 5mC detected in 2iL rapidly increased upon FA treatment in RAS^lox/lox^, ERF^KO^ and RAS^KO^; ERF^KO^ ESC. However, FA-RAS^KO^ ESC remained largely hypomethylated (fig. S7B). To get further insights into methylation dynamics on a genome-wide manner, we performed reduced representation bisulfite sequencing (RRBS) in all genotypes and pluripotent states (Fig. 7C, D) (*31*). 2iL-RAS^lox/lox^ ESC presented low levels of methylation (6.55% of analyzed CpG sites are methylated), which were increased after differentiation (21.75%). Furthermore, 2iL-ERF^KO^ ESC showed higher average methylation levels in 2iL (12.1%) and FA (24.7%) conditions, consistent with their bias toward differentiation and their less optimal naïve transcriptional network (Fig. 7C, D). Conversely, RAS^KO^ ESC showed extremely low methylation levels in 2iL conditions (1.55%) reaching similar levels to those found in 2iL-RAS^lox/lox^ ESC when differentiated (6.9%) (Fig. 7C, D). As expected, the defect in methylation observed in RAS^KO^ ESC was rescued upon ERF deletion (Fig. 7C, D). Interestingly, ERF^KO^ ESC showed higher level of methylation around transcription start sites as well as in CpG islands (Fig. 7E), especially after differentiation. Collectively, our data showed that ERF regulates negatively the expression of Lin28 and DNMT3 proteins. The altered expression of these proteins might underlie the global disbalance in the methylation levels at a genome-wide level in ERF-deficient cells.

## DISCUSSION

Preceding implantation, the cells residing within the naïve ICM of the blastocyst transition through a distinct phase of primed pluripotency in preparation for lineage commitment. This transition is initiated by fibroblast growth factor 4 (FGF4)-dependent activation of the RAS/MAPK pathway (*32, 33*). Despite the relevance of the MAPK pathway during this critical period, little is known about how RAS proteins instruct this transition. Here, we used RAS-deficient ESC for a careful dissection of the FGF pathway and demonstrated that the transcriptional factor ERF is the sole MAPK-dependent switch that controls the naïve-to-primed transition.

FA-RAS^KO^ ESC are characterized by an active naïve transcriptional network, high expression of OTX2 and elevated expression of primed markers. This suggests that 1) activation of the primed transcriptional program does not require full exit from naïve pluripotency, and 2) FGF/MAPK is the main pathway controlling the exit from naïve pluripotency. Indeed, our data revealed that ERF imposes an exquisite level of coordination during the transition from naïve to primed pluripotency. In the absence of MAPK signaling, chromatin bound ERF ensures elevated expression of naïve pluripotent factors, including NANOG (Fig. 4). High levels of NANOG maintain the undifferentiated state of ESC in the absence of LIF and might shield naïve embryonic cells from premature commitment (*24, 34*). Accordingly, ERF^KO^ ESC showed decreased levels of NANOG and undergo a partial exit from naïve pluripotency under 2iL conditions (Fig. 4). Thus, our data suggest that ERF supports naïve pluripotency, at least partially, by sustaining elevated levels of NANOG. ERF also controls negatively the expression of LIN28 proteins as 2iL-ERF^KO^ ESC showed increased levels of both LIN28A and B. Although LIN28 proteins regulate the expression of DNMT3 proteins by sequestering let-7 microRNAs, we did not detect upregulation of the methyltransferases in naïve ERF^KO^ ESC (Fig. 7). This is likely due to high levels of PRDM14, which represses expression of DNMT3 methyltransferases by recruiting Polycomb repressive complex 2 to their promoters, and it is regulated by ERF (Fig. 5E) (*35*). Thus, coordinated PRDM14 downregulation and increased LIN28 expression through ERF-dependent mechanisms could contribute to DNMT3 upregulation and increased methylation during the transition to primed pluripotency. Although DNMT3A and DNMT3B are dispensable for a successful exit from naïve pluripotency, *de novo* DNA methylation facilitates a timely progression towards a primed state (*35*). In fact, DNMT3A/B double knockout ESC exhibited a delayed exit from the naïve state and defective expression of the post-implantation markers OCT6, FGF5 or OTX2 (*36*). This is ultimately due to persistent NANOG expression after 2iL withdrawal as DNMT3A/B deficiency leads to reduced CpG methylation at the proximal NANOG promoter (*36, 37*). Similarly, FA-RAS^KO^ ESC showed negligible expression of DNMT3B and, therefore, high levels of NANOG. Interestingly, ERF^KO^ ESC showed increased DNA methylation compared to RAS^lox/lox^ ESC and is particularly evident in genes and CpG islands after the transition to EpiLSC (Fig. 7E). Naïve ERF^KO^ ESC have low levels of DNMT3 expression but elevated levels of DNMT3L, the catalytically inactive regulatory factor of *de novo* DNA methyltransferases (fig. S7C), which can contribute to this bias during differentiation (*38*). Furthermore, DPPA3, also known as Stella and implicated in preventing excessive DNA methylation by sequestering the E3 ubiquitin ligase UHRF1, is also decreased in ERF^KO^ ESC (Fig. S7C) (*39, 40*). Importantly, downregulation of DPPA3 mediated by DNMT3 methylation has been shown to be a key event in the naïve to primed conversion (*41*). Collectively, elevated levels of DNMT3L and low levels of DPPA3 in naïve conditions could contribute to the aberrant level of methylation detected in FA-ERF^KO^. Nevertheless, these changes in methylation do not result in major transcriptional changes in FA-ERF^KO^ compared to their respective control cells.

FA-RAS^KO^ ESC showed high expression of OTX2, which has been shown a critical transcription factor for thew maintenance of intermediate states of pluripotency (*7*). The consequence of having high levels of OTX2 in this context can be inferred by previous studies (*25*). Indeed, OTX2 over-expression has been shown to redirect OCT4 to previously inaccessible sites in EpiLSC, decrease expression of naïve markers (TBX3 or ESRRB), and induce EpiLSC genes, such as FGF5 (*25*). Interestingly, these effects take place in the presence of MEKi, a situation that is mirrored in FA-RAS^KO^ ESC. The ability of OTX2 to engage new enhancer sequences not only depends on its levels but also on the cooperative help of additional transcription factors (*25*). This cooperation might explain the enrichment of OTX2 in active naïve enhancers in FA-RAS^KO^ ESC. Moreover, our data also showed that, in addition to OCT4, NANOG is mobilized to sites that become active in EpiLSCs, which results in partial or full transcriptional activation (Fig. 6E). In this context, NANOG relocation might facilitate OTX2 positioning as it has been shown to promote chromatin accessibility and TF recruitment together with BRG1, part of the large remodeling complex SWI/SNF (*24*).

Finally, we showed that FA-RAS^KO^ ESC can be maintained in culture under primed conditions and could be a suitable genetic model to study intermediate states of pluripotency. Indeed, FA-RAS^KO^ ESC mirrored transcriptionally cells that have departed 12-24 hours from naïve pluripotency and are reminiscent of the recently described pluripotent intermediate states (*6–8*). Future studies will be necessary to determine the similarity of all these intermediate states with corresponding *in vivo* counterparts. Nevertheless, pluripotency can be considered as a dynamic property associated to different transient stem cell states that can be recapitulated by using different inhibitors and/or growth factors. By using RAS^KO^ ESC we focused specifically on the FGF pathway and the role of RAS proteins during the naïve to primed transition.

Future studies will be needed to determine if ERF plays a role in other cell fate decisions during early embryonic development such as in primitive endoderm specification, where FGF4-dependent activation of the MAPK pathway is also necessary. Moreover, the control of ERF over PRDM14 expression suggested that ERF might also play an important role in germline fate (*42*). In conclusion, here we demonstrated the essential role of ERF as regulator of the timely transition to primed pluripotency.

## MATERIALS AND METHODS

### Embryo Culture

All the animal work included here was performed in C57BL/6J mice obtained from the Jackson Laboratory in compliance with the NIH Animal Care & Use Committee (ACUC) Guideline for Breeding and Weaning. 4-weeks old female mice were injected intraperitoneally with 5IU Pregnant Mare Serum Gonadotropin (PMSG, Prospec) followed by 5 IU human Chorionic Gonadotropin (hCG, Sigma-Aldrich) 46-48 hours later. Alternatively, 8-weeks naturally pregnant females were euthanized, and embryos collected in M2 media (MR-015-D, Sigma-Aldrich) at indicated time points after hCG injection: E2.75, E3.5, E4.0, E4.75. The sex of embryos was not determined. Embryos were fixed in 4% Paraformaldehyde (Electron Microscopy Sciences) for 10 min, permeabilized for 30 min in 0.3% Triton X-100 and 0.1M Glycine in PBS 1X and blocked for 1 hour (1% BSA, 0.1% Tween in PBS 1X). Embryos were incubated overnight with primary antibodies (see STAR methods), washed in 0.1% Tween in PBS 1X and incubated with the secondary antibody accordingly for 1 hour at room temperature. Embryos were imaged using a Nikon Ti2-E microscope (Nikon Instruments) equipped with a CSU-W1 spinning disk (Yokogawa), Photometrics Prime BSI sCMOS (Photometrics), and 60x Nikon Apochromat TIRF objective (NA = 1.49). Z-stacks were acquired with a x-y pixel size of 0.11 mm and z-step of 0.9 mm. For quantification, embryo z-stack images were quantified using Imaris Bitplane (Oxford Instruments). 3D surfaces were rendered based on nuclear DAPI-staining and the corresponding regions were used to quantify the fluorescence intensity of ERF, NANOG, and KLF4. For experiment shown in Fig. 1A shows one representative experiment with the following embryos and cells used for quantification: Exp 1: E2.75: 3 Embryos, 24 cells in total. E3.5: 3 Embryos, 117 cells total, E4.0: 4 Embryos, 168 cells total, E4.75: 1 Embryo: 75 cells. Exp 2: E3.5: 8 embryos, E3.75: 4 embryos, E4.0: 6 embryos.

### Cell culture and differentiation

*N-Ras^-/-^; H-Ras^-/-^; K-Ras^f/f^; Ubiq-Cre^ERT2^* (RAS^lox/lox^) (*15*) ESC were grown in N2B27 media supplemented with 2i/LIF (1 mM PD0325901, 3 mM CHIR99021, both from Tocris and 1:500 LIF, made in house). N2B27 media consisted of a 1:1 mix of DMEM/F12 and Neurobasal Medium, 1X N2 supplement, 1X B27 supplement, 0.1 mM nonessential amino acids, 55 μM β-mercaptoethanol and 1% penicillin/streptomycin (all from Life Technologies). Cells were routinely cultured in 0.1% gelatinized plates and passaged with Accutase (Gibco) unless otherwise indicated. To induce EpiLSC differentiation, cells were grown for few passages in plates pretreated with 10 μg/ml polyL-ornithine and 5 μg/ml laminin (Corning). A total of 200,000-300,000 cells per 10 cm^2^ were plated on plates pretreated with 5 μg/ml Fibronectin (Millipore) in N2B27 media supplemented with 1% KOSR, 12 ng/ml FGF2 (R&D systems) and 20 ng/ml Activin A (PeproTech) for 48 hours including daily media changes. For inducing CRE-mediated deletion of K-RAS allele, we incubated RAS^lox/lox^ ESC with 1μM 4-hydroxytamoxifen (4-OHT, Sigma-Aldrich) for 6 days before performing any experiment. To maintain EpiLSC in culture, ESCs were plated in N2B27 media with 12 ng/ml FGF2, 20 ng/ml Activin A and 1 mM XAV939 on Fibronectin-coated plates at a density of 10,000 cells per cm^2^. EpiLSC were passaged the first time with Accutase including 1 mM Y27632 to enhance plating efficiency. Media was changed every other day and passaged every 2-3 days. HEK293T (American Type Culture Collection) cells were grown in DMEM, 10% FBS, and 1% penicillin/streptomycin.

To target a short half-life form of eGFP (deGFP) in the endogenous REX1 gene, we generated a targeting vector by inserting deGFP in pCR^®^-Blunt II TOPO^®^ (Zero Blunt TOPO PCR cloning kit, Invitrogen). Homology arms PCR-amplified from endogenous sequences upstream of the start codon and downstream of the stop codon from REX1 were cloned to flank pCR^®^-Blunt II TOPO^®^-deGFP. In addition, specific small guide RNA (sgRNA) sequences targeting the surroundings of the REX1 start codon were cloned into the plasmid pX330-U6-Chimeric_BB-CBh-hSpCas9 (Addgene, 42230) (*44*). The sequences of the sgRNAs were designed with the Genetic Perturbation Platform sgRNA designer tool (https://portals.broadinstitute.org/gpp/public/analysis-tools/sgrna-design). Both plasmids were transfected in ESC using Jetprime (Polyplus transfection) to induce the targeting and cell sorted based on GFP intensity to isolate individual clonal ESC lines. See Table S1 for primer information.

### Clonogenicity Assay

ESC were withdrawn of 2i/LIF for 48 hours and plated at single cell density (50 cells/cm^2^) in N2B27 media with 2i/LIF on plates coated with 0.1% gelatin (Sigma). At day 5, alkaline phosphatase staining was performed using the Alkaline Phosphatase Detection Kit (Millipore). Colonies were counted manually. At least three independent experiments with three replicates per experiment were performed.

### Generation of self-organizing embryonic spheres

ESC growing on gelatinized plates in N2B27 media with 2i/LIF were dissociated with Accutase and washed with PBS before their resuspension in growth factor reduced Matrigel (Corning) at a concentration of 10,000 cells/20μl of matrigel. The suspension was deposited in drops in 8-μwell Ibidi microplates and incubated at 37°C until the Matrigel solidified. Wells were then filled with N2B27 media without 2i/LIF and cultured for 48-72 hours at 37°C and 5% CO2. ESC-derived spheres were fixed in 4% PBS–paraformaldehyde (PFA) for 10 min at room temperature. Permeabilization was performed in PBS containing 0.3% Triton X-100 (Sigma) and 0.1 M glycine (Sigma) for 30 minutes at room temperature. Spheres were incubated with primary antibodies at 4°C overnight, followed by incubation with corresponding fluorescently conjugated Alexa Fluor secondary antibodies for 2 hours at room temperature. Both primary and secondary antibodies were diluted in PBS containing 1% BSA (Sigma) and 0.1% Tween20 (Sigma). See Table S2 for antibody information.

### Western blot

Cells were lysed in 50 mM Tris pH 8, 8 M Urea (Sigma) and 1% Chaps (Millipore) followed by 30 min of shaking at 4°C. 20 μg of supernatants were run on 4%-12% NuPage Bis-Tris Gel (Invitrogen) and transferred onto Nitrocellulose Blotting Membrane (GE Healthcare). Membranes were blocked in 5% skim milk (Millipore) and 0.1% Tween 20 (Sigma) in PBS. Membranes were incubated with the primary antibody overnight at 4°C (see STAR methods), followed by incubation with HRP-conjugated secondary antibodies (1:5000) for 1 h at room temperature. Membranes were developed using SuperSignal West Pico PLUS (Thermo Scientific). See Table S2 for antibody information.

### Dot blots

Trypsinized ESC were lysed for 2-3 hours at 55°C with lysis buffer (100 mM Tris-HCl pH 8, 5 mM EDTA, 0.2% SDS, 20 mM NaCl and 100 μg/ml Proteinase K). DNA was isolated by adding an equal volume of Phenol/Chloroform/Isoamyl to the samples and using phase-lock tubes (5PRIME Phase Lock Gel™ Heavy, Quantabio), followed by an extraction with identical volume of Chloroform. DNA was precipitated with 2 volumes of 100% Ethanol plus 0.3 M NaAc, washed with 70% Ethanol and resuspended in water. A total of 500 ng of DNA was diluted in 0.3 M NaOH and denatured at 42°C for 12 minutes. After incubation, the samples were rapidly transferred by spotting each sample into a nitrocellulose membrane (Nitrocellulose Blotting Membrane, GE Healthcare). After the transfer, DNA was crosslinked with a Stratalinker^®^ UV crosslinker (Stratagene) using the Autocrosslink setting. The membrane was blocked in 5% skim milk (Millipore) and 0.1% Tween 20 (Sigma) in PBS, incubated with 1:500 dilution of the anti-5mC antibody (see STAR methods) overnight at 4°C, followed by incubation with HRP-conjugated secondary antibody (1:5000) for 1 h at room temperature. Membrane was developed using SuperSignal West Pico PLUS (Thermo Scientific). See Table S2 for antibody information.

### Flow cytometry

For flow cytometry experiments, cells were dissociated into single cell suspensions and analyzed for GFP gene expression using a FACS Fortessa (BD Biosciences). DAPI was added to detect cells with compromised membrane integrity. Data was analyzed using FlowJo. At least two independent experiments were performed.

### Immunofluorescence

Cells were fixed in 4 % Paraformaldehyde (Electron Microscopy Sciences) for 10 min at room temperature, permeabilized in 100 mM Tris-HCl pH 7.4, 50 mM EDTA pH 8.0, 0.5% Triton X-100 and incubated with the corresponding primary antibodies overnight (see STAR methods). This was followed by incubation with corresponding fluorescently conjugated Alexa Fluor secondary antibodies for 2 h at room temperature. Both primary and secondary antibodies were diluted in PBS containing 1% BSA (Sigma) and 0.1% Tween20 (Sigma). Images were acquired using either a Nikon spinning disk confocal microscope (CSU-W1) or a Zeiss LSM880 Airyscan microscope. See Table S2 for antibody information.

### RNAseq and data analysis

RNA was isolated using the ISOLATE II RNA Mini Kit (Bioline) following manufacturer’s recommendations. DNA libraries for RNAseq analysis were prepared using NEBNext Ultra II Directional RNA Library Prep Kit for Illumina (New England Biolabs, NEB) and NEBNext rRNA Depletion Kit (Human/Mouse/Rat) (NEB) according to the manufacturer’s protocol. Sequencing was performed on the Illumina NextSeq550 (75bp pair-end reads). RNAseq reads were adapter trimmed using fastp v.0.20.0 (*45*). Transcript expression was quantified via mapping to mouse gencode v25 transcripts using salmon (*46*). Identification of differentially expressed genes between samples was performed using DESEQ2 (PMID: 25516281). RNAseq coverage tracks were generated by aligning RNAseq reads to UCSC version mm10 of the mouse genome using the STAR v2.6.1a aligner (*47*) followed by application of the ‘bamCoverage’ utility from deeptools (*48*) to generate signal track files with the following parameters: normalization=RPKM, bin_size=50, smooth_length=1. For comparison to other published RNAseq data sets, gene counts across samples were quantile-normalized using the limma package (*49*). Batch correction was then performed on quantile-normalized counts using COMBAT (*50*). See Table S3 for further software and algorithm information.

### CUT&RUN protocol

The CUT&RUN protocol was slightly modified from (*51, 52*). In brief, cells were washed thrice with Wash Buffer (20 mM HEPES-KOH pH 7.5, 150 mM NaCl, 0.5 mM spermidine, Roche complete Protease Inhibitor tablet EDTA free) and bound to activated Concanavalin A beads (Polysciences) for 10 minutes at room temperature. Cells were then permeabilized in Digitonin Buffer (0.05 % Digitonin and 0.1% BSA in Wash Buffer) and incubated with corresponding antibodies at 4°C for 2 hours. For negative controls, Guinea Pig anti-Rabbit IgG (Antibodies-online) was used. Following antibody incubation, cells were washed with Digitonin Buffer and incubated with a hybrid protein A-protein G-Micrococcal nuclease (pAG-MNase) at 4°C for 1 hour. Samples were washed in Digitonin Buffer, resuspended in 150 μl Digitonin Buffer and equilibrated to 0°C on ice water for 5 minutes. To initiate MNase cleavage, 3 μl 100 mM CaCl2 was added to cells and after 1 hour of digestion, reactions were stopped with the addition of 150 μl 2x Stop Buffer (340 mM NaCl, 20 mM EDTA, 4 mM EGTA, 0.02 % Digitonin, 50 μg/ml RNase A, 50 μg/ml Glycogen). Samples were incubated at 37°C for 10 minutes to release DNA fragments and centrifuged at 16,000 g for 5 minutes. Supernatants were collected and a mix of 1.5 μl 20% SDS / 2.25 μl 20 mg/ml Proteinase K was added to each sample and incubated at 65°C for 35 minutes. DNA was precipitated with ethanol and sodium acetate and pelleted by high-speed centrifugation at 4°C, washed, air-dried and resuspended in 10 μ 0.1x TE. See Table S2 for antibody information.

### Library preparation and sequencing

The entire precipitated DNA obtained from CUT&RUN was used to prepare Illumina compatible sequencing libraries. In brief, end-repair was performed in 50 μl of T4 ligase reaction buffer, 0.4 mM dNTPs, 3 U of T4 DNA polymerase (NEB), 9 U of T4 Polynucleotide Kinase (NEB) and 1 U of Klenow fragment (NEB) at 20°C for 30 minutes. End-repair reaction was cleaned using AMPure XP beads (Beckman Coulter) and eluted in 16.5 μl of Elution Buffer (10 mM Tris-HCl pH 8.5) followed by A-tailing reaction in 20 μl of dA-Tailing reaction buffer (NEB) with 2.5 U of Klenow fragment exo-(NEB) at 37°C for 30 minutes. The 20 μl of the A-tailing reaction were mixed with Quick Ligase buffer 2X (NEB), 3000 U of Quick Ligase (NEB) and 10 nM of annealed adaptor (Illumina truncated adaptor) in a volume of 50 μl and incubated at room temperature for 20 min. The adaptor was prepared by annealing the following HPLC-purified oligos: 5’-Phos/GATCGGAAGAGCACACGTCT-3’ and 5’-ACACTCTTTCCCTACACGACGCTCTTCCGATC*T-3’ (*phosphorothioate bond). Ligation was stopped by adding 50 mM of EDTA, cleaned with AMPure XP beads and eluted in 14 μl of Elution Buffer. All volume was used for PCR amplification in a 50 μl reaction with 1 μM primers TruSeq barcoded primer p7, 5’-CAAGCAGAAGACGGCATACGAGATXXXXXXXXGTGACTGGAGTTCAGACGTGTGCTCTTCCGATC*T-3’ and TruSeq barcoded primer p5 5’-AATGATACGGCGACCACCGAGATCTACACXXXXXXXXACACTCTTTCCCTACACGACGCTCTTCCGATC*T-3’ (* represents a phosphothiorate bond and XXXXXXXX a barcode index sequence), and 2X Kapa HiFi HotStart Ready mix (Kapa Biosciences). The temperature settings during the PCR amplification were 45 s at 98°C followed by 15 cycles of 15 s at 98°C, 30 s at 63°C, 30 s at 72°C and a final 5 min extension at 72°C. PCR reactions were cleaned with AMPure XP beads (Beckman Coulter), run on a 2% agarose gel and a band of 300bp approximately was cut and gel purified using QIAquick Gel Extraction Kit (QIAGEN). Library concentration was determined with KAPA Library Quantification Kit for Illumina Platforms (Kapa Biosystems). Sequencing was performed on the Illumina NextSeq550 (75bp pair-end reads).

### Cut&Run data processing

Data were processed using a modified version of Cut&RunTools (*53*). Reads were adapter trimmed using fastp v.0.20.0 (*45*). An additional trimming step was performed to remove up to 6bp adapter from each read. Next, reads were aligned to the mm10 genome using bowtie2 (*54*) with the ‘dovetail’ and ‘sensitive’ settings enabled. macs2 (*55*) was used to call peaks with q-value cutoff < 0.01. Normalized (RPKM) signal tracks were generated using the ‘bamCoverage’ utility from deepTools with parameters bin-size=25, smooth length=75, and ‘center_reads’ and ‘extend_reads’ options enabled (*48*). See Table S3 for further software and algorithm information.

### Processing for published ChIP datasets

External next generation sequencing data were downloaded from the Sequence Read Archive (SRA) and analyzed as follows. These analyses include a re-analysis of our original datasets (*9*). Reads were aligned to the mm10 genome using bowtie2 (*54*). Duplicate reads were removed using MarkDuplicates from the Picard toolkit (“Picard Toolkit.” 2019. Broad Institute, GitHub Repository. http://broadinstitute.github.io/picard/). Normalized (RPKM) signal tracks were generated bamCoverage utility from deepTools (*48*), using the parameters bin-size=25, smooth length=75, ‘center_reads’ and ‘extend_reads’. For paired-end data, read mates were extended to the fragment size defined by the two read mates. For single-end ChIP-seq data, reads were extended to the estimated fragment length estimated by phantompeakqualtools (*56*). See Table S3 for further software and algorithm information.

### Identification of OCT4 binding sites in ESC and EpiLSC

Fastq files from published OCT4 ChIP-seq data for ESC (SRR1202455, SRR1202456), EpiLSC plus and minus Activin-A (SRR1202468, SRR1202469), and associated input controls (SRR1202465, SRR1202464, SRR1202477 SRR1202478) (*25*) were downloaded from the Sequence Read Archive (SRA). Single-end reads were aligned to the mm10 genome using bwa (*57*). Duplicate reads were removed using MarkDuplicates from the Picard toolkit (“Picard Toolkit.” 2019. Broad Institute, GitHub Repository. http://broadinstitute.github.io/picard/), and peaks for each sample were called using macs2 (*55*) with q-value cutoff < 0.01 and extension length determined using phantompeakqualtools (*56*). Diffbind (*58*) using the DeSeq2 method was used to determine differentially bound peaks, treating EpiLSC plus and minus Activin-A samples as replicate experiments as was done in the original study (*25*). Peaks were determined to be ESC-or EpiLC-specific if they differed by 2-fold read concentration with p-val<0.01 and FDR <0.03. A subset of peaks with > mean read concentration for EpiLC and ESC with <0.5 fold difference were selected as “common” or shared peaks. See Table S3 for further software and algorithm information.

### RRBS data processing

DNA libraries for RRBS analysis were prepared using the Premium RRBS kit (Diagenode) following manufacturer’s recommendations. Sequencing was performed on the Illumina NextSeq550 (75bp single-end reads). Single-end RRBS reads were adapter and quality trimmed (phred33 score>=20) using trimgalore v0.6.5 (http://www.bioinformatics.babraham.ac.uk/projects/trim_galore) with the RRBS option invoked. Bismark v0.22.1 (*59*) was used to align reads to UCSC version mm10 of the mouse genome. CpG methylation was extracted using bismark ignoring the first 4 bases of the read after inspection of m-bias plots. Differentially methylated regions were called using the methylKit package (*60*). See Table S3 for further software and algorithm information.

## QUANTIFICATION AND STATISTICAL ANALYSIS

All information regarding statistical details of the experiments, number of experiments, statistical tests used, number of cells/experiments can be found in the corresponding figure legends o methods section.

## Acknowledgments

We thank Bechara Saykali for critical reading of the manuscript, Christa Buecker for sharing reagents, Mariano Barbacid for sharing and Sagrario Ortega for isolating RAS^lox/lox^ ESC. David Goldstein and the CCR Genomics Core for sequencing support and Ferenc Livak and the CCR Flow cytometry Core for experimental support. Research in S.R. laboratory is supported by the Intramural Research Program of the NIH. T.O. is supported by a Helen Hay Whitney fellowship.

## Author contributions

M.V-S. and S.R. conceived the study. M.V-S., T.O., designed and performed experiments. C.N.D. and M.F. provided technical support. D.T. and P.C.F. analyzed sequencing data. A.D.T. and M.J.K. analyzed confocal microscopy data. S.R. supervised the study and wrote the manuscript with comments from all authors.

## Data and materials availability

All data needed to evaluate the conclusions in the paper are present in the paper and/or the Supplementary Materials. The RNA-seq and Cut&Run data generated in this study have been deposited in the GEO database under accession GSE162581. Additional data related to this paper may be requested from the authors.

## REFERENCES

1. J. Wray, T. Kalkan, A.G. Smith, The ground state of pluripotency. Biochem. Soc. Trans. 38, 1027–32 (2010).

2. J. Nichols, A. Smith, A. Naive and primed pluripotent states. Cell Stem Cell 4, 487–492 (2009).

3. L. Weinberger, M. Ayyash, N. Novershtern, J.H. Hanna, Dynamic stem cell states: naive to primed pluripotency in rodents and humans. Nat. Rev. Mol. Cell Biol. 17, 155–69 (2016).

4. Q.L. Ying, J. Wray, J. Nichols, L. Batlle-Morera, B. Doble, J. Woodgett, P. Cohen, A. Smith, The ground state of embryonic stem cell self-renewal. Nature 453, 519–23 (2008).

5. P.J. Tesar, J.G. Chenoweth, F.A. Brook, T.J. Davies, E.P. Evans, D.L. Mack, R.L., Gardner, R.D. McKay, New cell lines from mouse epiblast share defining features with human embryonic stem cells. Nature 448, 196–199 (2007).

6. A. Neagu, E. van Genderen, I. Escudero, L. Verwegen, D. Kurek, J. Lehmann, J. Stel, R.A.M. Dirks, G. van Mierlo, A. Maas, C. Eleveld, Y. Ge, A.T. den Dekker, R.W.W. Brouwer, W.F.J. van IJcken, M. Modic, M., Drukker, J.H. Jansen, N.C. Rivron, E.B. Baart, H. Marks, D. Ten Berge, In vitro capture and characterization of embryonic rosette-stage pluripotency between naive and primed states. Nat. Cell Biol. 22, 534–545 (2020).

7. M. Kinoshita, M., Barber, W. Mansfield, Y. Cui, D. Spindlow, G.G. Stirparo, S. Dietmann, J. Nichols, A. Smith, Capture of Mouse and Human Stem Cells with Features of Formative Pluripotency. Cell Stem Cell 28: DOI:https://doi.org/10.1016/j.stem.2020.11.005, (2020).

8. L. Yu, Y. Wei, H. Sun, A.K. Mahdi, C.P. Arteaga, M. Sakurai, D.A. Schmitz, C. Zheng, E.D. Ballard, J. Li, N. Tanaka, A. Kohara, D. Okamura, A.A. Mutto, Y. Gu, P.J. Ross, J. Wu, Derivation of Intermediate Pluripotent Stem Cells Amenable to Primordial Germ Cell Specification. Cell Stem Cell 28: DOI:https://doi.org/10.1016/j.stem.2020.11.003 (2020).

9. C. Mayor-Ruiz, T. Olbrich, M. Drosten, E. Lecona, M. Vega-Sendino, S. Ortega, O. Dominguez, M. Barbacid, S. Ruiz O. Fernández-Capetillo, ERF deletion rescues RAS deficiency in mouse embryonic stem cells. Genes&Dev 32, 568–576 (2018).

10. D.N. Sgouras, M.A. Athanasiou, G.J. Jr. Beal, R.J. Fisher, D.G. Blair, G.J. Mavrothalassitis, ERF: an ETS domain protein with strong transcriptional repressor activity, can suppress ets-associated tumorigenesis and is regulated by phosphorylation during cell cycle and mitogenic stimulation. EMBO J. 14, 4781–4793 (1995).

11. L. Le Gallic, D. Sgouras, G. Jr. Beal, G. Mavrothalassitis, Transcriptional repressor ERF is a Ras/mitogen-activated protein kinase target that regulates cellular proliferation. Mol. Cell. Biol. 19, 4121–4133 (1999).

12. L. Le Gallic, L., Virgilio, P. Cohen, B. Biteau, G. Mavrothalassitis, ERF nuclear shuttling, a continuous monitor of Erk activity that links it to cell cycle progression. Mol. Cell. Biol. 24, 1206–18 (2004).

13. E. Vorgia, A. Zaragkoulias, I. Peraki, G. Mavrothalassitis, Suppression of Fgf2 by ETS2 repressor factor (ERF) is required for chorionic trophoblast differentiation. Mol. Reprod. Dev. 84, 286–295 (2017).

14. S. Nowotschin, M. Setty, Y.Y. Kuo, V. Liu, V., Garg, R. Sharma, C.S. Simon, N. Saiz, R. Gardner, S.C. Boutet, D.M. Church, P.A. Hoodless, A.K. Hadjantonakis, D. Pe’er, The emergent landscape of the mouse gut endoderm at single-cell resolution. Nature 569, 361–367 (2019).

15. M. Drosten, A. Dhawahir, E.Y. Sum, J. Urosevic, C.G. Lechuga, L.M. Esteban, E. Castellano, C. Guerra, E. Santos, M. Barbacid, Genetic analysis of Ras signalling pathways in cell proliferation, migration and survival. EMBO J. 29, 1091–1104 (2010).

16. K. Hayashi, H. Ohta, K. Kurimoto, S. Aramaki, M. Saitou Reconstitution of the mouse germ cell specification pathway in culture by pluripotent stem cells. Cell 146, 519–532 (2011).

17. F. Nakaki, K. Hayashi, H. Ohta, K. Kurimoto, Y. Yabuta, M. Saitou, Induction of mouse germ-cell fate by transcription factors in vitro. Nature 501, 222–226. (2013).

18. P. Yang, S.J. Humphrey, S. Cinghu, R. Pathania, A.J. Oldfield, D. Kumar, D. Perera, J.Y.H. Yang, D.E. James, M. Mann, R. Jothi, Multi-omic Profiling Reveals Dynamics of the Phased Progression of Pluripotency. Cell Syst. 8, 427–445 (2019).

19. I. Bedzhov, M. Zernicka-Goetz, Self-organizing properties of mouse pluripotent cells initiate morphogenesis upon implantation. Cell 156, 1032–1044 (2014).

20. M.N. Shahbazi, A. Scialdone, N. Skorupska, A. Weberling, G. Recher, M. Zhu, A. Jedrusik, L.G. Devito, L. Noli, I.C. Macaulay, C. Buecker, Y. Khalaf, D. Ilic, T. Voet, J.C. Marioni, M. Zernicka-Goetz Pluripotent state transitions coordinate morphogenesis in mouse and human embryos. Nature 552: 239–243 (2017).

21. T. Kalkan, N. Olova, M. Roode, C. Mulas, H.J. Lee, I. Nett, H. Marks, R. Walker, H.G. Stunnenberg, K.S. Lilley, J. Nichols, W. Reik, P. Bertone, A. Smith, Tracking the embryonic stem cell transition from ground state pluripotency. Development 144, 1221–1234 (2017).

22. D. Hnisz, B.J. Abraham, T.I. Lee, A. Lau, V. Saint-André, A.A. Sigova, H.A. Hoke, R.A. Young, Super-enhancers in the control of cell identity and disease. Cell 155, 934–947 (2013).

23. W.A. Whyte, D.A. Orlando, D. Hnisz, B.J. Abraham, C.Y. Lin, M.H. Kagey, P.B. Rahl, T.I. Lee, R.A. Young, Master transcription factors and mediator establish super-enhancers at key cell identity genes. Cell 153, 307–319 (2013).

24. V. Heurtier, N. Owens, I. Gonzalez, F. Mueller, C. Proux, D. Mornico, P. Clerc, A. Dubois, P. Navarro, The molecular logic of Nanog-induced self-renewal in mouse embryonic stem cells. Nat. Commun. 10, 1109 (2019).

25. C. Buecker, R. Srinivasan, Z. Wu, E. Calo, D. Acampora, T. Faial, A. Simeone, M. Tan, T. Swigut, J. Wysocka, Reorganization of enhancer patterns in transition from naive to primed pluripotency. Cell Stem Cell 14, 838–53 (2014).

26. T. Zhang, Z. Zhang, Q. Dong, J. Xiong, B. Zhu, Histone H3K27 acetylation is dispensable for enhancer activity in mouse embryonic stem cells. Genome Biol. 21, 45 (2020).

27. J. Zhang, S. Ratanasirintrawoot, S. Chandrasekaran, Z. Wu, S.B. Ficarro, C. Yu, C.A. Ross, D. Cacchiarelli, Q. Xia, M. Seligson, G. Shinoda, W. Xie, P. Cahan, L. Wang, S.C. Ng. S. Tintara, C. Trapnell, C. Onder, Y.H. Loh, T. Mikkelsen, P. Sliz, M.A. Teitell, J.M. Asara, J.A. Marto, H. Li, J.J. Collins, G.Q. Daley, LIN28 Regulates Stem Cell Metabolism and Conversion to Primed Pluripotency. Cell Stem Cell 19, 66–80 (2016).

28. J. Balzeau, M.R. Menezes, S. Cao, J.P. Hagan, The LIN28/let-7 Pathway in Cancer. Front. Genet. 8, 31 (2017).

29. G. Auclair, S. Guibert, A. Bender, M. Weber, Ontogeny of CpG island methylation and specificity of DNMT3 methyltransferases during embryonic development in the mouse. Genome Biol. 15, 545 (2014).

30. S. Takahashi, S., Kobayashi, I. Hiratani Epigenetic differences between naïve and primed pluripotent stem cells. Cell Mol. Life Sci. 75, 1191–1203 (2018)

31. A. Meissner, A., Gnirke, G.W. Bell, B. Ramsahoye, E.S. Lander, R. Jaenisch, Reduced representation bisulfite sequencing for comparative high-resolution DNA methylation analysis. Nucleic Acids Res. 33, 5868–5877 (2005).

32. M. Kang, A. Piliszek, J. Artus, A.K. Hadjantonakis, FGF4 is required for lineage restriction and salt-and-pepper distribution of primitive endoderm factors but not their initial expression in the mouse. Development 140: 267–279 (2013).

33. Y. Yamanaka, F. Lanner, J. Rossant, FGF signal-dependent segregation of primitive endoderm and epiblast in the mouse blastocyst. Development 137: 715–724 (2010).

34. K. Mitsui, Y. Tokuzawa, H. Itoh, K. Segawa, M. Murakami, K. Takahashi, M. Maruyama, M., Maeda, S. Yamanaka, The homeoprotein Nanog is required for maintenance of pluripotency in mouse epiblast and ES cells. Cell 113, 631–642 (2003).

35. M. Yamaji, J. Ueda, K. Hayashi, H. Ohta, Y. Yabuta, K. Kurimoto, R. Nakato, Y. Yamada, K. Shirahige, M. Saitou, PRDM14 ensures naive pluripotency through dual regulation of signaling and epigenetic pathways in mouse embryonic stem cells. Cell Stem Cell 12, 368–382 (2013).

36. M.A. Li, P.P. Amaral, P. Cheung, J.H. Bergmann, M. Kinoshita, T. Kalkan, M. Ralser, S. Robson, F. von Meyenn, M. Paramor, F. Yang, C. Chen, J. Nichols, D.L. Spector, T. Kouzarides, L. He, A. Smith, A lncRNA fine tunes the dynamics of a cell state transition involving Lin28, let-7 and de novo DNA methylation. Elife 6, e23468 (2017).

37. C.R. Farthing, G. Ficz, R.K. Ng, C.F. Chan, S. Andrews, W. Dean, M. Hemberger, W. Reik, Global mapping of DNA methylation in mouse promoters reveals epigenetic reprogramming of pluripotency genes. PLoS Genet. 4, e1000116 (2008).

38. N. Veland, Y. Lu, S. Hardikar, S., Gaddis, Y. Zeng, B. Liu, M.R. Estecio, Y. Takata, K. Lin, M.W. Tomida, J. Shen, D. Saha, H. Gowher, H., Zhao, T. Chen, DNMT3L facilitates DNA methylation partly by maintaining DNMT3A stability in mouse embryonic stem cells. Nucleic Acids Res. 47, 152–167. (2019).

39. W. Du, Q. Dong, Z. Zhang, B. Liu, T. Zhou, R.M. Xu, H. Wang, B. Zhu, Y. Li, Stella protein facilitates DNA demethylation by disrupting the chromatin association of the RING finger-type E3 ubiquitin ligase UHRF1. J Biol Chem. 294, 8907–8917 (2019).

40. T. Nakamura, Y. Arai, H. Umehara, M. Masuhara, T. Kimura, H. Taniguchi, T. Sekimoto, M. Ikawa, Y. Yoneda, M. Okabe, S. Tanaka, K. Shiota, T. Nakano, PGC7/Stella protects against DNA demethylation in early embryogenesis. Nat. Cell Biol. 9, 64–71 (2007).

41. H. Sang, D. Wang, S. Zhao, J. Zhang, Y. Zhang, J. Xu, X. Chen, Y. Nie, K. Zhang, S. Zhang, Y. Wang, N. Wang, F. Ma, L. Shuai, Z. Li, N. Liu, Dppa3 is critical for Lin28a-regulated ES cells naïve-primed state conversion. J. Mol. Cell Biol. 11, 474–488 (2019).

42. N. Grabole, J. Tischler, J.A. Hackett, S. Kim, F. Tang, H.G. Leitch, E. Magnúsdóttir, M.A. Surani, Prdm14 promotes germline fate and naive pluripotency by repressing FGF signaling and DNA methylation. EMBO Rep. 14, 629–637. (2013).

43. C. Galonska, M.J. Ziller, R. Karnik, A. Meissner, Ground State Conditions Induce Rapid Reorganization of Core Pluripotency Factor Binding before Global Epigenetic Reprogramming. Cell Stem Cell 17, 462–470 (2015).

44. L. Cong, F.A. Ran, D. Cox, S. Lin, R. Barretto, N. Habib, P.D. Hsu, X. Wu, W. Jiang, L.A. Marraffini, F. Zhang, Multiplex genome engineering using CRISPR/Cas systems. Science, 339: 819–23 (2013)

45. S. Chen, Y. Zhou, Y., Chen, J. Gu, fastp: an ultra-fast all-in-one FASTQ preprocessor. Bioinformatics 34: 884–i890 (2018)

46. R. Patro, G. Duggal, M.I. Love, R.A. Irizarry, C. Kingsford, Salmon provides fast and bias-aware quantification of transcript expression. Nat. Methods 14, 417–419 (2017).

47. A. Dobin, C.A. Davis, F. Schlesinger, J. Drenkow, C. Zaleski, S. Jha, P. Batut, M. Chaisson, T.R. Gingeras, STAR: ultrafast universal RNA-seq aligner. Bioinformatics 29: 15–21. (2013).

48. F. Ramírez, D.P. Ryan, B. Grüning, V. Bhardwaj, F. Kilpert, A.S. Richter, S. Heyne, F. Dündar, T. Manke, deepTools2: a next generation web server for deep-sequencing data analysis. Nucleic Acids Res 44: 160–165 (2016)

49. M.E. Ritchie, B. Phipson, D. Wu, Y. Hu, C.W. Law, W. Shi, G.K. Smyth, limma powers differential expression analyses for RNA-sequencing and microarray studies. Nucleic Acids Res. 43: e47 (2015).

50. W.E. Johnson, C. Li, A. Rabinovic, Adjusting batch effects in microarray expression data using empirical Bayes methods. Biostatistics 8: 118–127 (2007).

51. P.J. Skene, S. Henikoff, An efficient targeted nuclease strategy for high-resolution mapping of DNA binding sites. Elife 6, e21856 (2017).

52. M.P. Meers, T.D. Bryson, J.G. Henikoff, S. Henikoff, Improved CUT&RUN chromatin profiling tools. Elife 8, e46314 (2019).

53. Q. Zhu, N. Liu, S.H. Orkin, G.C. Yuan, CUT&RUNTools: a flexible pipeline for CUT&RUN processing and footprint analysis. Genome Biol. 20: 192 (2019).

54. B. Langmead, S.L. Salzberg, Fast gapped-read alignment with Bowtie 2. Nat. Methods 9: 357–359. (2012).

55. Y. Zhang, T. Liu, C.A. Meyer, J. Eeckhoute, D.S. Johnson, B.E. Bernstein, C. Nusbaum, R.M. Myers, M. Brown, W. Li, X.S. Liu, Model-based analysis of ChIP-Seq (MACS). Genome Biol. 9: R137. (2008).

56. P.V. Kharchenko, M.Y. Tolstorukov, P.J. Park, Design and analysis of ChIP-seq experiments for DNA-binding proteins. Nat. Biotechnol. 26: 1351–1359 (2008).

57. H. Li, R. Durbin, Fast and accurate short read alignment with Burrows-Wheeler transform. Bioinformatics 25, 1754–1760 (2009).

58. R. Stark G.D. Brown DiffBind: Differential Binding Analysis of ChIP-Seq Peak Data. Bioconductor. Available online at: http://bioconductor.org/packages/release/bioc/html/DiffBind.html. (2011).

59. F. Krueger, S.R. Andrews, Bismark: a flexible aligner and methylation caller for Bisulfite-Seq applications. Bioinformatics 27: 1571–1572 (2011).

60. A. Akalin, M. Kormaksson, S. Li, F.E. Garrett-Bakelman, M.E. Figueroa, A. Melnick, C.E. Mason, methylKit: a comprehensive R package for the analysis of genome-wide DNA methylation profiles. Genome Biol. 13, R87 (2012).

61. D. van Dijk, R. Sharma, J. Nainys, K. Yim, P. Kathail, A.J. Carr, C. Burdziak, K.R. Moon, C.L. Chaffer, D. Pattabiraman, B. Bierie, L. Mazutis, G. Wolf, S. Krishnaswamy, D. Pe’er, Recovering Gene Interactions from Single-Cell Data Using Data Diffusion. Cell 174: 716–729.e27 (2018).

62. J. Wu, B. Huang, H. Chen, Q. Yin, Y. Liu, Y. Xiang, B. Zhang, B., Liu, Q. Wang, W. Xia, W. Li, Y. Li, J. Ma, X. Peng, H. Zheng, J. Ming, W. Zhang, J. Zhang, G. Tian, F. Xu, Z. Chang, J. Na, X. Yang, W. Xie, The landscape of accessible chromatin in mammalian preimplantation embryos. Nature 534, 652–657 (2016).

