## Supplementary Information for "The ETS Transcription Factor ERF controls the exit from the naïve pluripotent state"

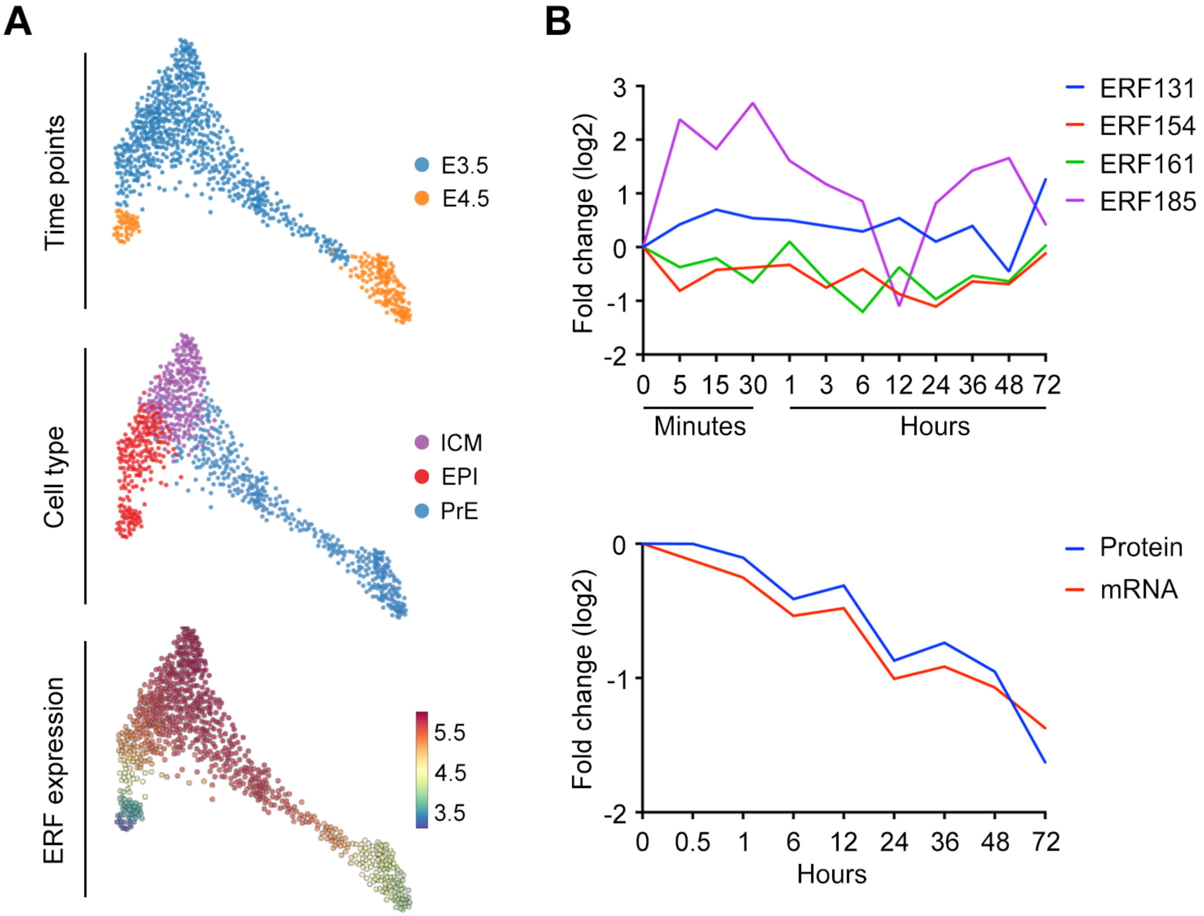
A
Time points
E3.5
E4.5
Cell type
ICM
EPI
PrE
ERF expression
5.5
4.5
3.5
B
Fold change (log2)
3
2
1
0
-1
-2
0 5 15 30 1 3 6 12 24 36 48 72
Minutes
Hours
ERF131
ERF154
ERF161
ERF185
Fold change (log2)
0
-1
-2
0 0.5 1 6 12 24 36 48 72
Hours
Protein
mRNA

**Figure S1**

**Figure S1: ERF expression decreases upon exit from naïve pluripotency.** (**A**) Forced-directed layouts of E3.5 and E4.5 single cell RNAseq data showing the temporal relation between ICM (inner cell mass), EPI (epiblast) and PrE (primitive endoderm). Cells are colored by timepoint (upper panel) or cell type (middle panel). Data was obtained from (*14*). In the lower panel, ERF gene expression is shown. Cells are colored by gene expression post-imputation with the MAGIC algorithm (*14, 61*). Plots were obtained from https://endoderm-explorer.com/. (**B**) Temporal dynamics of relative ERF phosphorylation levels (upper panel) and ERF protein and mRNA levels (lower panel) compared to timepoint 0 hours during the transition from ESC to EpiLSC. Data was obtained from (Yang et al., 2019).

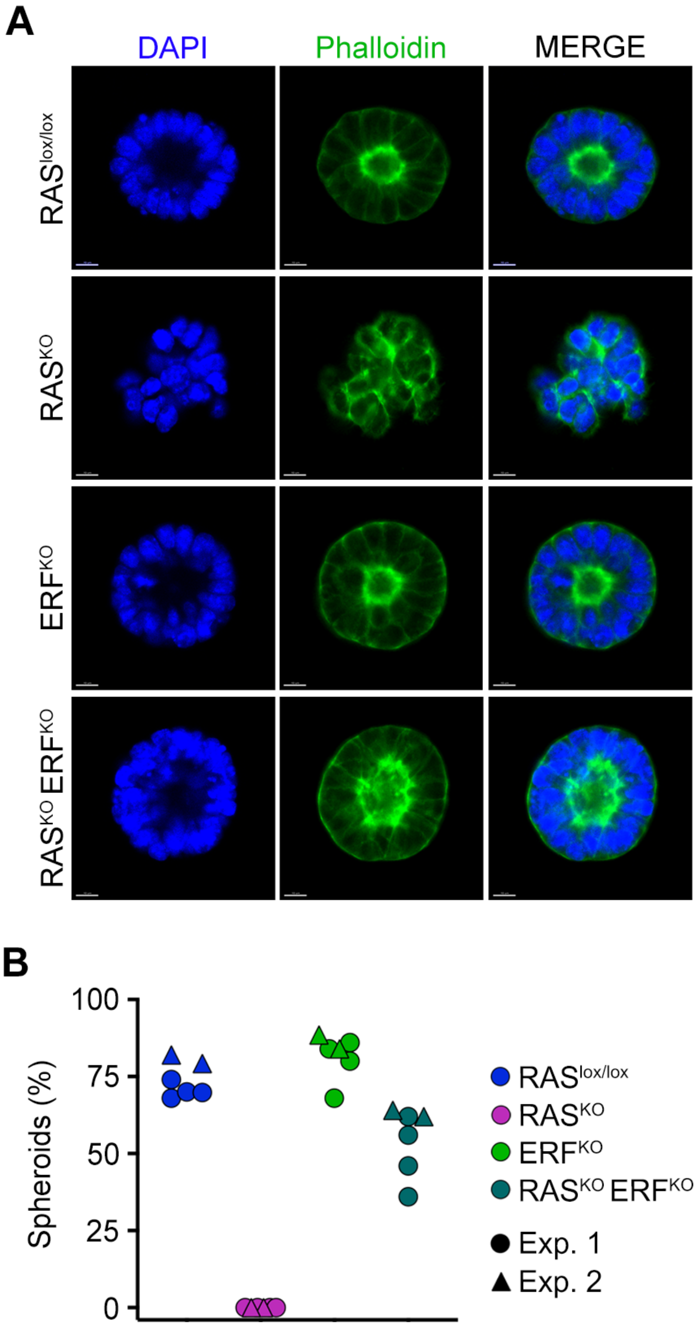
A
DAPI
Phalloidin
MERGE
RASlox/lox
RASKO
ERFKO
RASKO ERFKO
B
100
75
50
25
0
Spheroids (%)
RASlox/lox
RASKO
ERFKO
RASKO ERFKO
Exp. 1
Exp. 2

**Figure S2**

**Figure S2: Downregulation of ERF is necessary for successful exit from naïve pluripotency**. (**A**) Central confocal optical sections of $RAS^{lox/lox}$, $ERF^{KO}$, $RAS^{KO}$, and $RAS^{KO}$; $ERF^{KO}$ embryonic cell rosettes embedded in matrigel 48 hours after seeding and stained with phalloidin (green). DAPI was used to visualize nuclei. Scale bars, 10μm. (**B**) Graph showing the percentage of morphologically organized embryonic rosettes in all genotypes based on the staining from (A). Two independent experiments are shown and at least, a total of 50 rosettes were counted.

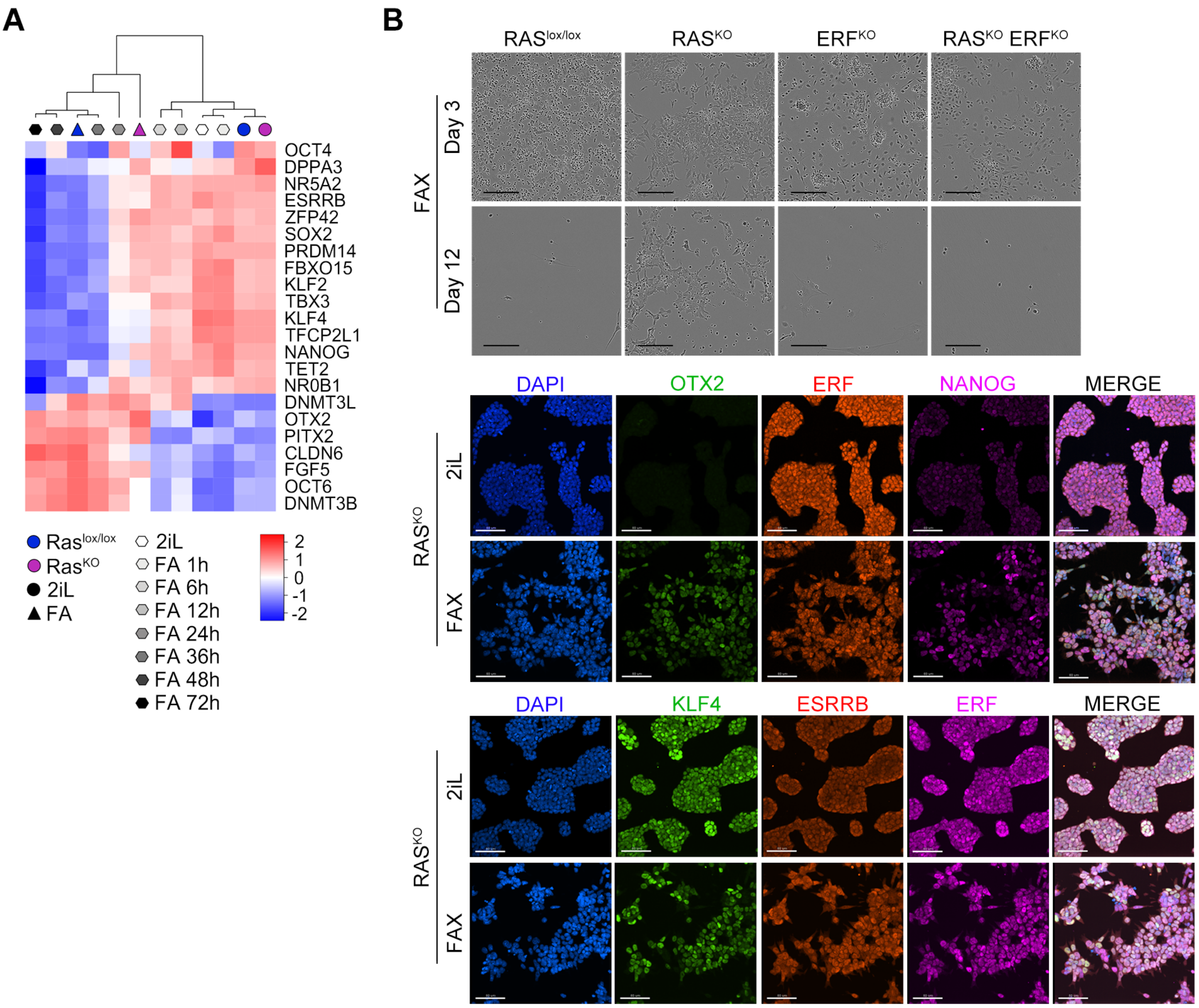
A
OCT4
DPPA3
NR5A2
ESRRB
ZFP42
SOX2
PRDM14
FBXO15
KLF2
TBX3
KLF4
TFCP2L1
NANOG
TET2
NR0B1
DNMT3L
OTX2
PITX2
CLDN6
FGF5
OCT6
DNMT3B
Ras lox/lox
Ras KO
2iL
FA
2iL
FA 1h
FA 6h
FA 12h
FA 24h
FA 36h
FA 48h
FA 72h
2
1
0
-1
-2
B
RAS lox/lox
RAS KO
ERF KO
RAS KO ERF KO
FAX
Day 3
Day 12
DAPI
OTX2
ERF
NANOG
MERGE
RAS KO
2iL
FAX
DAPI
KLF4
ESRRB
ERF
MERGE
RAS KO
2iL
FAX

**Figure S3**

**Figure S3: ERF controls the transition to primed pluripotency.** (**A**) Heatmap generated from RNAseq datasets showing 2iL and FA treated $RAS^{lox/lox}$ and $RAS^{KO}$ ESC along with RNAseq datasets from a time course experiment during EpiLSC induction (0, 1, 6, 12, 24, 36, 48 and 72 hours) (*18*). Shown are the averaged values of at least 2 replicates. (**B**) Bright field images of $RAS^{lox/lox}$, $ERF^{KO}$, $RAS^{KO}$, and $RAS^{KO}$; $ERF^{KO}$ ESC cultures in EpiLSC media (FAX) 3 and 12 days after the media switch (upper panels). Note that only EpiLSC cultures were able to be maintained with $RAS^{KO}$ ESC. Scale bars, 200μm. In the lower panel, immunofluorescence analysis of 2iL and FAX treated $RAS^{KO}$ ESC and stained for OTX2 (green), ERF (red) and NANOG (purple) are shown. DAPI was used to visualize nuclei. Scale bars, 80μm.

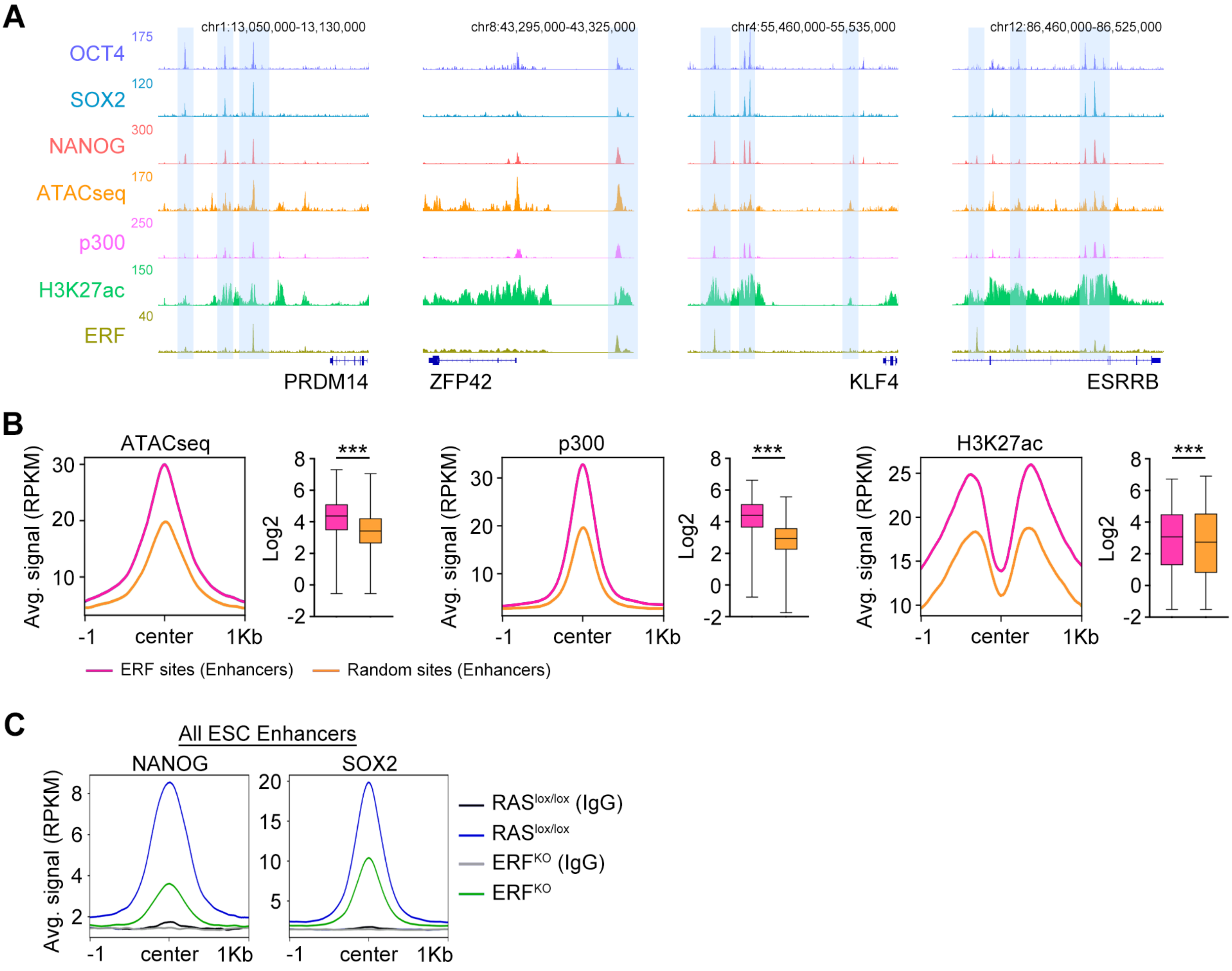
A
chr1:13,050,000-13,130,000
chr8:43,295,000-43,325,000
chr4:55,460,000-55,535,000
chr12:86,460,000-86,525,000
OCT4
175
SOX2
120
NANOG
300
ATACseq
170
p300
250
H3K27ac
150
ERF
40
PRDM14
ZFP42
KLF4
ESRRB
B
ATACseq
p300
H3K27ac
Avg. signal (RPKM)
Log2
***
-1
center
1Kb
ERF sites (Enhancers)
Random sites (Enhancers)
C
All ESC Enhancers
NANOG
SOX2
RASlox/lox (IgG)
RASlox/lox
ERFKO (IgG)
ERFKO

**Figure S4**

**Figure S4: Chromatin-bound ERF ensures an optimal naïve pluripotency state.** (**A**) Genome browser tracks showing OCT4, SOX2, NANOG (*43*), H3K27ac, P300 (*25*), ATACseq (*62*) and ERF (Mayor-Ruiz et al., 2018) normalized read count at PRDM14, ESRRB, KLF4 and REX1 (ZFP42) naïve associated genes in ESC. ERF binding sites are highlighted. (**B**) ChIPseq read density plot showing H3K27ac, p300 (*25*), as well as ATACseq signal (*62*) at 2074 ERF-binding sites at enhancers (pink) or 2074 randomly selected non-ERF bound enhancers (orange). *** = $p<0.001$, T-student. (**C**) Cut&Run read density plot showing SOX2 and NANOG occupancy in all ESC enhancers (10672) in $RAS^{lox/lox}$ (blue) and $ERF^{KO}$ (green) ESC cultured in 2iL. Corresponding inputs (IgG) are also shown as reference control.

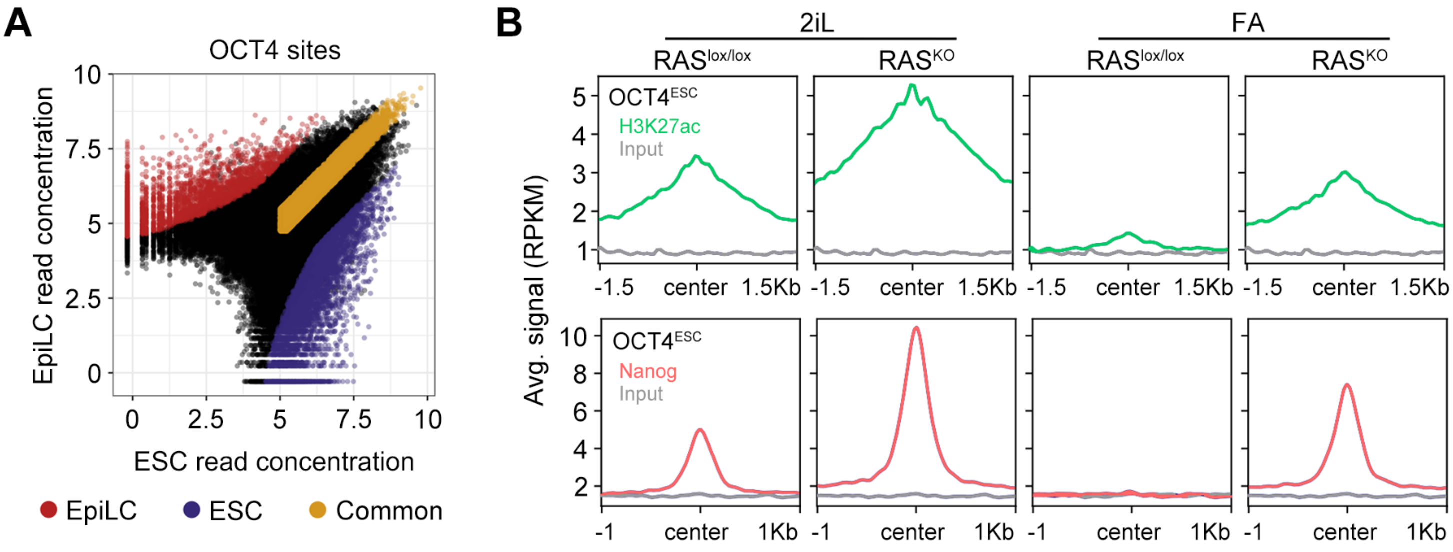
A
OCT4 sites
EpiLC read concentration
ESC read concentration
10
7.5
5
2.5
0
EpiLC
ESC
Common
B
2iL
FA
RASlox/lox
RASKO
OCT4ESC
H3K27ac
Input
Nanog
Avg. signal (RPKM)
-1.5
center
1.5Kb
-1
1Kb

Figure S5

**Figure S5: The naïve enhancer landscape is active in rosette-like FA-RAS$^{KO}$ ESC.** (**A**) Classification of OCT4 sites based on change in occupancy during differentiation. Replicates of OCT4 peaks in ESC and EpiLC (*25*) were used to determine differentially bound peaks using DiffBind (*58*). A threshold of p-val<0.01 and FDR<0.03 and <2-fold read concentration was chosen to define ESC-specific sites (blue) and EpiLC-specific sites (red). A set of least-changed OCT4 peaks between was treated as common or shared sites (orange). (**B**) Cut&Run read density plot showing H3K27ac (plots above, green) and NANOG (plots below, red) occupancy in all OCT4$^{ESC}$ sites in RAS$^{lox/lox}$ and RAS$^{KO}$ cultured in 2iL or differentiated to EpiLSC (FA). Corresponding inputs (IgG) are also shown as reference control.

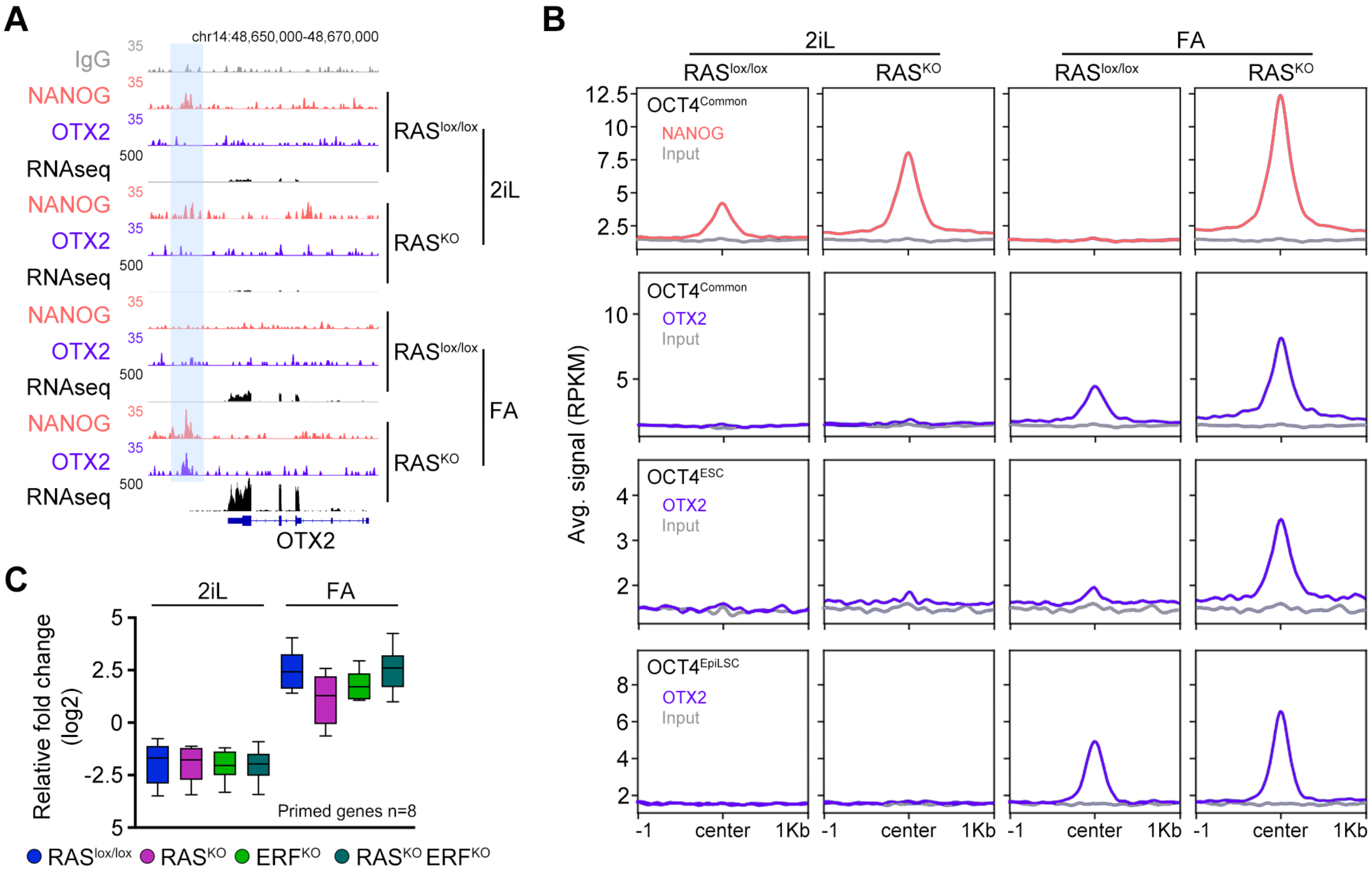

A
chr14:48,650,000-48,670,000
IgG
NANOG
OTX2
RNAseq
RAS^lox/lox
RAS^KO
2iL
FA
OTX2
B
2iL
FA
RAS^lox/lox
RAS^KO
OCT4^Common
NANOG
Input
OTX2
OCT4^ESC
OCT4^EpiLSC
Avg. signal (RPKM)
-1
center
1Kb
C
2iL
FA
Relative fold change (log2)
Primed genes n=8
RAS^lox/lox
RAS^KO
ERF^KO
RAS^KO ERF^KO

**Figure S6**

**Figure S6: OTX2 co-occupy binding sites with NANOG in FA-RAS$^{KO}$ ESC.** (**A**) Genome browser tracks showing NANOG and OTX2 occupancy at the OTX2 gene and RNAseq RPKM read count in RAS$^{lox/lox}$ and RAS$^{KO}$ ESC cultured in 2iL or differentiated to EpiLSC (FA). Inputs (IgG) are also shown as a reference control. Blue squares showed ERF binding sites. (**B**) Cut&Run read density plot showing NANOG (red) and OTX2 (purple) occupancy in the indicated OCT4$^{Common}$, OCT4$^{ESC}$ and OCT4$^{EpiLSC}$ sites in RAS$^{lox/lox}$ and RAS$^{KO}$ ESC cultured in 2iL or differentiated to EpiLSC (FA). Corresponding inputs (IgG) are also shown as reference control. (**C**) Graph showing the relative fold change (log2) expression of a subset of 8 genes associated to primed pluripotency (*Dnmt3a*, *Dnmt3b*, *Fgf5*, *Fgf15*, *OCT6*, *Wnt8a*, *Otx2* and *Dnmt3l*) in the different genotypes in 2iL or differentiated to EpiLSC (FA). For each gene, data was normalized to the average across all samples.

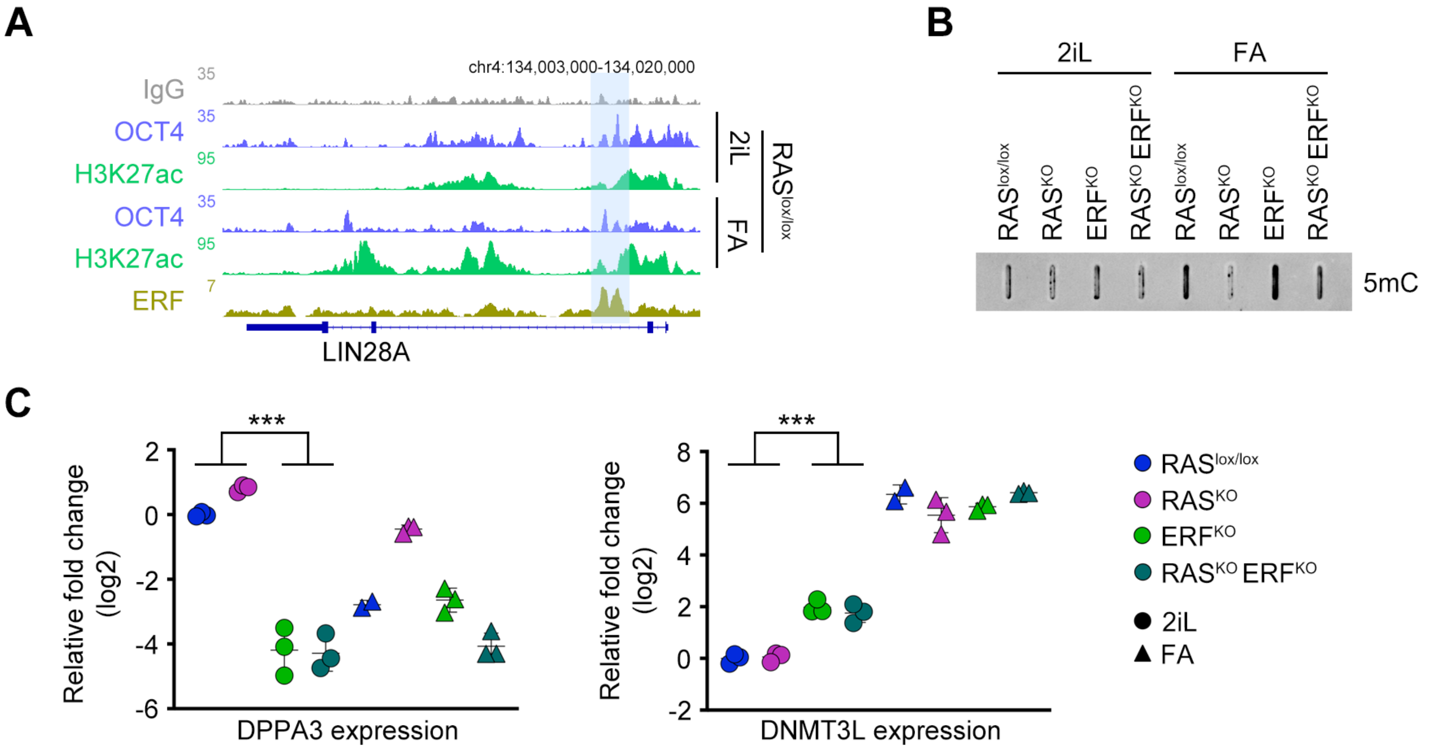
A
chr4:134,003,000-134,020,000
IgG
OCT4
H3K27ac
OCT4
H3K27ac
ERF
35
35
95
35
95
7
2iL
FA
RASlox/lox
LIN28A
B
2iL
FA
RASlox/lox
RASKO
ERFKO
RASKO ERFKO
RASlox/lox
RASKO
ERFKO
RASKO ERFKO
5mC
C
***
Relative fold change (log2)
DPPA3 expression
***
Relative fold change (log2)
DNMT3L expression
RASlox/lox
RASKO
ERFKO
RASKO ERFKO
2iL
FA

Figure S7

**Figure S7: ERF controls methylation during naïve-to-primed transition.** (**A**) Genome browser tracks showing H3K27ac deposition and OCT4 occupancy at the LIN28A gene in $RAS^{lox/lox}$ ESC cultured in 2iL or differentiated to EpiLSC (FA). ERF binding profile in $RAS^{KO}$ ESC is also shown. Inputs (IgG) are also shown as a reference control. Blue squares showed ERF binding sites. (**B**) Dot blot analysis to detect the levels of 5mC in ESC from all genotypes cultured in 2iL or differentiated to EpiLSC (FA). (**C**) Plots showing the relative fold change (log2) expression for DPPA3 (left panel) and DNMT3L (right panel) in ESC from all genotypes cultured in 2iL or differentiated to EpiLSC (FA). Data is shown as triplicates. For each gene, data was normalized to the average across all samples. *** = $p<0.001$, T-student.

**Table S1:** Primers used in this study.

| Oligonucleotides |
|---|
| Primer for left homology REX1 arm:<br>5'-REX1-KpnI-F: ACGTGGTACCTCTTTGCCTTACAGAGAAGCC |
| Primer for left homology REX1 arm:<br>5'-REX1-SacI-R: ACGTGAGCTCGTTGTCTTAGCTGCTTCCTTC |
| Primer for right homology REX1 arm:<br>3'-REX1-NotI-F: ACGTGCGGCCGCAGGTGGAGACAGATTGTCCTC |
| Primer for right homology REX1 arm:<br>3'-REX1-XhoI-F: ACGTCTCGAGTTGCCTTAAGTTCTGTATGC |
| eGFPd2-F: ACAACATGGTGAGCAAGGGCGAGGAGC |
| eGFPd2-R: ACGTCTACACATTGATCCTAGCAGAAG |
| sgRNA-REX1-F1: CACCGAGTGGCCAGAAAGGGCCGGG |
| sgRNA-REX1-R1: AAACCCCGGCCCTTTCTGGCCACTC |
| sgRNA-REX1-F2: CACCGCCATATCCGCATCCACACCG |
| sgRNA-REX1-R2: AAACCGGTGTGGATGCGGATATGGC |

**Table S2:** Antibodies used in this study.

| ANTIBODIES | SOURCE | IDENTIFIER |
|---|---|---|
| ERF Antibody (E-9) | Santa Cruz Biotechnology | Cat# sc-398269 |
| Anti-Nanog antibody | Abcam | Cat# ab80892, RRID:AB_2150114 |
| Anti Nanog (Mouse) pAb | Cosmo Bio USA | Cat# REC-RCAB002P-F |
| Mouse KLF4 Antibody | R&D systems | Cat# AF3158, RRID:AB_2130245 |
| Mouse Podocalyxin Antibody | R&D systems | Cat# MAB1556, RRID:AB_2166010 |
| Human Otx2 Antibody | R&D systems | Cat# AF1979, RRID:AB_2157172 |
| Anti-Otx1 + Otx2 antibody | Abcam | Cat# ab21990, RRID:AB_776930 |
| Alexa Fluor™ 488 Phalloidin | Invitrogen | Cat# A12379 |
| Anti-OCT6 Antibody, clone KT110 | Millipore | Cat# MABN738, RRID:AB_2876862 |
| Human ERR beta/NR3B2 Antibody | R&D systems | Cat# PP-H6705-00, RRID:AB_2100412 |
| Anti-Sox2 Antibody | Millipore | Cat# AB5603, RRID:AB_2286686 |
| Anti-Histone H3 (acetyl K27) antibody | Abcam | Cat# ab4729, RRID:AB_2118291 |
| p44/42 MAPK (Erk1/2) Antibody | Cell Signaling Technology | Cat# 9102, RRID:AB_330744 |
| Phospho-p44/42 MAPK (Erk1/2) (Thr202/Tyr204) Antibody | Cell Signaling Technology | Cat# 9101, RRID:AB_331646 |
| LIN28A (D1A1A) XP® Rabbit mAb | Cell Signaling Technology | Cat# 8641, RRID:AB_10997528 |
| LIN28B Antibody | Cell Signaling Technology | Cat# 5422, RRID:AB_10697489 |
| Monoclonal Anti-α-Tubulin antibody | Sigma-Aldrich | Cat# T9026, RRID:AB_477593 |
| Anti-Dnmt3b antibody | Abcam | Cat# ab122932, RRID:AB_10933207 |
| DNMT3A Antibody | Novus Biological | Cat# NB120-13888, RRID:AB_789607 |
| Pan-RAS (Ab-3) Mouse mAb (RAS 10) | Millipore | Cat# OP40-100UG, RRID:AB_213400 |
| Anti-5-methylcytosine (5-mC) antibody [33D3] | Abcam | Cat# ab10805, RRID:AB_442823 |
| Guinea Pig anti-Rabbit IgG (Heavy & Light Chain) Antibody | Antibodies-Online | Cat# ABIN101961, RRID:AB_10775589 |
| Goat anti-Rabbit IgG (H+L) Secondary Antibody, HRP | Thermo Fisher Scientific | Cat# 31466, RRID:AB_10960844 |
| Goat anti-Mouse IgG (H+L) Secondary Antibody, HRP | Thermo Fisher Scientific | Cat# 31431, RRID:AB_10960845 |
| Chicken anti-Rabbit IgG (H+L) Cross-Adsorbed Secondary Antibody, Alexa Fluor 488 | Thermo Fisher Scientific | Cat# A-21441, RRID:AB_2535859 |

| | | |
|---|---|---|
| Goat anti-Mouse IgG (H+L) Cross-Adsorbed Secondary Antibody, Alexa Fluor 568 | Thermo Fisher Scientific | Cat# A-11004, RRID:AB_2534072 |
| Donkey anti-Mouse IgG (H+L) Highly Cross-Adsorbed Secondary Antibody, Alexa Fluor 568 | Thermo Fisher Scientific | Cat# A10037, RRID:AB_2534013 |
| Chicken anti-Goat IgG (H+L) Cross-Adsorbed Secondary Antibody, Alexa Fluor 488 | Thermo Fisher Scientific | Cat# A-21467, RRID:AB_2535870 |
| Chicken anti-Rabbit IgG (H+L) Cross-Adsorbed Secondary Antibody, Alexa Fluor 647 | Thermo Fisher Scientific | Cat# A-21443, RRID:AB_2535861 |
| Donkey anti-Rabbit IgG (H+L) Highly Cross-Adsorbed Secondary Antibody, Alexa Fluor 568 | Thermo Fisher Scientific | Cat# A10042, RRID:AB_2534017 |
| Chicken anti-Rat IgG (H+L) Cross-Adsorbed Secondary Antibody, Alexa Fluor 488 | Thermo Fisher Scientific | Cat# A-21470, RRID:AB_2535873 |
| Chicken anti-Mouse IgG (H+L) Cross-Adsorbed Secondary Antibody, Alexa Fluor 647 | Thermo Fisher Scientific | Cat# A-21463, RRID:AB_2535869 |

**Table S3:** Software and Algorithms used in this study.

| Software and Algorithms | | |
|---|---|---|
| Cut&RunTools | Zhu et al., 2019 | https://bitbucket.org/gzhudfci/cutruntools/ |
| fastp v.0.20.0 | Chen et al., 2018 | https://github.com/OpenGene/fastp |
| bowtie2 | Langmead et al., 2012 | http://bowtie-bio.sourceforge.net/bowtie2/index.shtml |
| macs2 | Zhang et al., 2008 | https://github.com/macs3-project/MACS |
| deepTools | Ramirez et al., 2016 | https://github.com/deeptools/deepTools |
| Picard toolkit | | http://broadinstitute.github.io/picard/ |
| phantompeakqualtools | Kharchenko et al., 2008 | https://github.com/kundajelab/phantompeakqualtools |
| Diffbind v3.0.5 | Stark R. and Brown G. D. (2011). | https://bioconductor.org/packages/release/bioc/html/DiffBind.html |
| trimgalore v0.6.5 | | http://www.bioinformatics.babraham.ac.uk/projects/trim_galore |
| Bismark v0.22.1 | Krueger et al., 2011 | https://github.com/FelixKrueger/Bismark |
| methylKit v1.14.2 | Akalin et al., 2012 | https://bioconductor.org/packages/release/bioc/html/methylKit.html |
| Prism 8 | GraphPad | https://www.graphpad.com/ |
| FlowJo (10.1) | FlowJo LLC | https://www.flowjo.com/ |
| DNAnexus | DNAnexus | https://www.dnanexus.com/ |
| IGV | Robinson et al., 2011 | https://igv.org/ |
| R (3.5 and 4.0) | | www.r-project.org |
| GREAT | McLean et al., 2010 | http://great.stanford.edu/public/html/ |
| DESEQ2 | Love et al., 2014 | http://bioconductor.org/packages/release/bioc/html/DESeq2.html |
| Imaris Bitplane | Oxford Instruments | https://imaris.oxinst.com/ |
